# Whole-Genome Profiling of Ethyl Methanesulfonate Mutagenesis in Tomato

**DOI:** 10.1101/2022.04.19.488728

**Authors:** Prateek Gupta, Pankaj Singh Dholaniya, Kunnappady Princy, Athira Sethu Madhavan, Yellamaraju Sreelakshmi, Rameshwar Sharma

## Abstract

The induced mutations accelerate crop improvement by providing novel disease resistance and yield alleles. However, the alleles with no perceptible phenotype but having an altered function remain hidden in mutagenized plants. The whole-genome sequencing (WGS) of mutagenized individuals uncovers the complete spectrum of mutations in the genome. We sequenced 132 doubly ethyl methanesulfonate (EMS)-mutagenized lines of tomato and detected ca. 41 million SNPs and 5.5 million short-INDELs. We found a very high average density of mutations 1/3.05 Kb compared to other species. About 97% of the genome had mutations, including the genes, promoters, UTRs, and introns. More than 1/3^rd^ of genes in the mutagenized population had one or more deleterious mutations predicted by SIFT. Nearly 1/4^th^ of deleterious genes mapped on tomato metabolic pathways modulating multiple pathway steps. Contrary to the reported GC>AT transition bias for EMS, we found EMS also produced nearly equal AT>GC transitions. Comparing mutation frequency among synonymous codons revealed that the most preferred codon is least mutagenic towards EMS. The reduction in carotenoids in ζ-carotene isomerase mutant fruits and chloroplasts relocation loss in phototropin1 mutant validated the mutation discovery pipeline. Our study makes a large repertoire of mutations accessible to genetic studies and the breeding of tomato.

## Introduction

The impending climate changes and burgeoning human population have placed a marked emphasis on doubling global food production by 2050. Parallel to cereals that provide the most calorific values, there is an impetus to increase the yield and nutraceuticals in vegetable crops. Tomato, an important globally grown crop, is enriched in several nutraceuticals that are bereft in cereals. Like other crops, in tomato, the domestication pressure led to the loss in nutritional value, flavor, and disease resistance due to genetic erosion (**Tanksley and McCouch, 1997; Bauchet and Causse, 2012; Tieman et al., 2017**). The genome resequencing of a large number of tomato cultivars along with several wild relatives have highlighted the extent of loss of genetic diversity in modern tomato cultivars during domestication (**Aflitos et al., 2014; Lin et al., 2014; Gupta et al., 2020a**)

The genetic diversity of a domesticated crop can be augmented by introgression of chromosomal segments from wild relatives or by *de novo* induction of diversity by induced mutagenesis (**Kulus, 2018**). Before the advent of the genomics era, the diversity induced by mutagenesis was harnessed by the visual selection of the mutants displaying desired traits and backcrossing to the parent. The availability of genome sequences of crop plants expanded the scope of introgression of induced mutations by providing molecular markers (**Foolad, 2007; Simko et al., 2021**). It also facilitated functional genomic analysis of genes mutated by chemical/radiation mutagens, t-DNA, or transposon insertion.

Compared to induced mutagenesis, the t-DNA or transposon-mediated disruption of gene function remained limited to species easily amenable to the transformation, such as rice and Arabidopsis (**Ram et al., 2019; Provart et al., 2016**). In contrast, induced mutagenesis is not species-specific, but being random, it creates a large number of mutations across the whole genome (**Leitao, 2011**). The development of reverse genetic tools, particularly TILLING (Targeting Induced Local Lesions in Genomes) based on PCR screening, rendered it possible to identify the mutation in any gene (**McCallum et al., 2000**). The TILLING was not limited to detecting induced mutations; it was also applied to detect the SNPs present in natural accessions (**Comai et al., 2004**). The fast detection of mutations in any gene immensely widened the scope of the induced mutagenesis in crop plants.

To exploit the potential of TILLING, in several crop plants, such as rice, wheat, tomato, maize, mutant populations were generated (**Jacob et al., 2018**). In tomato, EMS-mutagenized populations for TILLING were made for several cultivars such as M82 (**Menda et al., 2004; Piron et al., 2010**), Red Setter (**Minoia et al., 2010**), TPAADASU (**Gady et al., 2009**), Micro-Tom (**Okabe et al., 2011**) and Arka Vikas (**Sharma et al., 2021**). An advantage of TILLING was that the pooled genomic DNA could be scanned for mutations in the target gene. Notwithstanding the convenience, TILLING is laborious and slow, as scanning of mutation at best could be done for 1.0-1.5 Kb genomic DNA, with no possibility of multiplexing. In addition, the precise identification of the mutated base needed Sanger Sequencing.

The above drawback was obviated by the next generation sequencing (NGS), where the mutagenized population could be analyzed using “TILLING by sequencing.” Herein the multi-dimensionally pooled genomic DNA was subjected to PCR to amplify a given gene. The PCR products were then pooled, followed by sequencing using NGS. The output data was analyzed to reveal rare induced mutations. The NGS-based TILLING was used for rice and wheat (**Tsai et al., 2011**), tomato (**Rigola et al., 2009, Gupta et al., 2017**), poplar (**Marroni et al., 2011**), peanut (**Guo et al., 2015**), and soybean (**Tsuda et al., 2015**). Though NGS speeded up the identification of mutations compared to conventional TILLING, yet the scope of NGS-based TILLING remained limited. The identification of a heterozygous mutation was limited to 64X pooled DNA. Most software were not robust enough to identify the mutations from the background noise barring a few such as CAMBA and GATK **(Gupta et al., 2017**).

The main drawback of TILLING was that mutations in only a few genes could be analyzed in one cycle, though the mutant lines bore mutations across the genome. The best approach is to identify the mutations across the genome by resequencing the genome. The reduction in whole-genome sequencing (WGS) cost made this approach feasible. However, the very large genomes of crop plants such as maize and higher ploidy levels of wheat made WGS an expensive approach. Presuming that most mutations in intergenic regions and introns do not elicit a phenotype, the sequencing was selectively done for the exome for maize and wheat. The whole-exome sequencing (WES) efficiently identified mutations in rice (a small genome plant) and wheat (a large genome plant) (**Henry et al., 2014**). In tetraploid and hexaploid wheat cultivars, the WES of 2,735 mutagenized lines identified more than 10 million mutations (**Krasileva et al., 2017**). Similarly, the WES of pollen-mutagenized maize plants identified nearly 0.2 million mutations in 1086 M_1_ lines (**Lu et al., 2018**).

Though the WES is more cost-effective than WGS, it has an inherent cost of capture probe designing and determining its efficiency. In maize, the efficiency of exome capture probes was about 83% (**Lu et al., 2018**). In addition, WES omits the significant number of mutations present in the intergenic regions, promoters, and introns (**Belkadi et al., 2015**). In tomato ‘Micro-Tom’ using exome capture, 241,391 mutations were identified in 95 M_2_ lines (**Yano et al., 2019**). While the mutations in the genic region are the ones that affect the traits, emerging evidences have indicated that mutations in introns, promoters, and intergenic region can also influence the trait. In tomato, a point mutation in the promoter of the *1-aminocyclopropane carboxylase2* gene reduced ethylene emission from the fruits and considerably prolonged the shelf life (**Sharma et al., 2021**). Thus, the potential of a mutagenized population can be best unlocked by the WGS, as it provides a repertoire of mutations across the genome.

In this study, we present the WGS of 132 EMS-mutagenized lines of tomato. Our analysis reveals that in addition to GC>AT transitions, EMS also caused a high frequency of AT> GC transitions. We have gene-indexed 41 million SNPs and 5.5 million INDELs in an open-access database called ITGV (Induced Tomato Genomic Variations, http://psd.uohyd.ac.in/itgv/). The ITGV allows users to search for mutations in the desired gene and visualize the mutations’ nature, functional effects, and protein alignment. The above collection of mutant lines provides the scientific community a genome-wide resource for tomato mutations. The mutant collection can be used for functional genomics of tomato and by breeders for trait improvement.

## RESULTS

### Whole-genome sequencing of EMS-mutagenized lines

To generate a genome-wide resource for tomato mutants, we did whole-genome sequencing (WGS) of 132 independent M_4_ EMS-mutagenized lines, including their progenitor cultivar, Arka Vikas (AV). In an earlier study (**Gupta et al., 2017**), we demonstrated the efficacy of the above-mutagenized population, where we sequenced 55 amplicons belonging to 25 genes derived from a 3-D pooled genomic DNA of 768 M_2_M_2_ EMS-mutagenized plants. The high-throughput WGS was carried out on the Illumina HiSeqX platform using 2X150 bp pair-end sequencing chemistry. The final output generated 4.72 terabytes of raw gene sequence data with 31.5 billion reads. On average, 238 million pair-end reads (35.75 Gb) were obtained for each M_4_ line with an average sequencing depth of 37.64-fold. The raw reads were filtered (Q30) using fastp software (**Chen et al., 2018**), and on average, 235 million (*range* 196-504 million) high-quality pair-end reads were obtained for each line after filtering with an average sequencing depth of 36.5-fold (**Table S1**, *range* 30.37-78.21 fold).

### The mutant population has a high frequency of A/T-to-G/C transitions

After variant filtering to remove artifacts, 46.5 million variations were detected, comprising ca. 41 million SNPs and 5.5 million short-INDELs in the analyzed M_4_ population. Contrary to the expectation, the G/C-to-A/T transitions were much lower, with an average of 27.86%. Surprisingly, the A/T-to-G/C transitions, which are expected to have a very low frequency, were 27.62%. Nearly the same pattern was reflected for conversion of individual nucleotides (C➔T, G➔A, A➔G, and T➔C) (**Figure 1A**). The remaining variations comprising transversion (GC> TA, AT> TA, AT> GC, GC> CG) ranged from 6% to 13%, with C>G and G>C transversion being the lowest (**Figure 1A,B**). This low frequency of C>G and G>C transversion is consistent with other studies, such as EMS-mutagenized rice lines (**Yan et al., 2021**).

**Figure 1.**
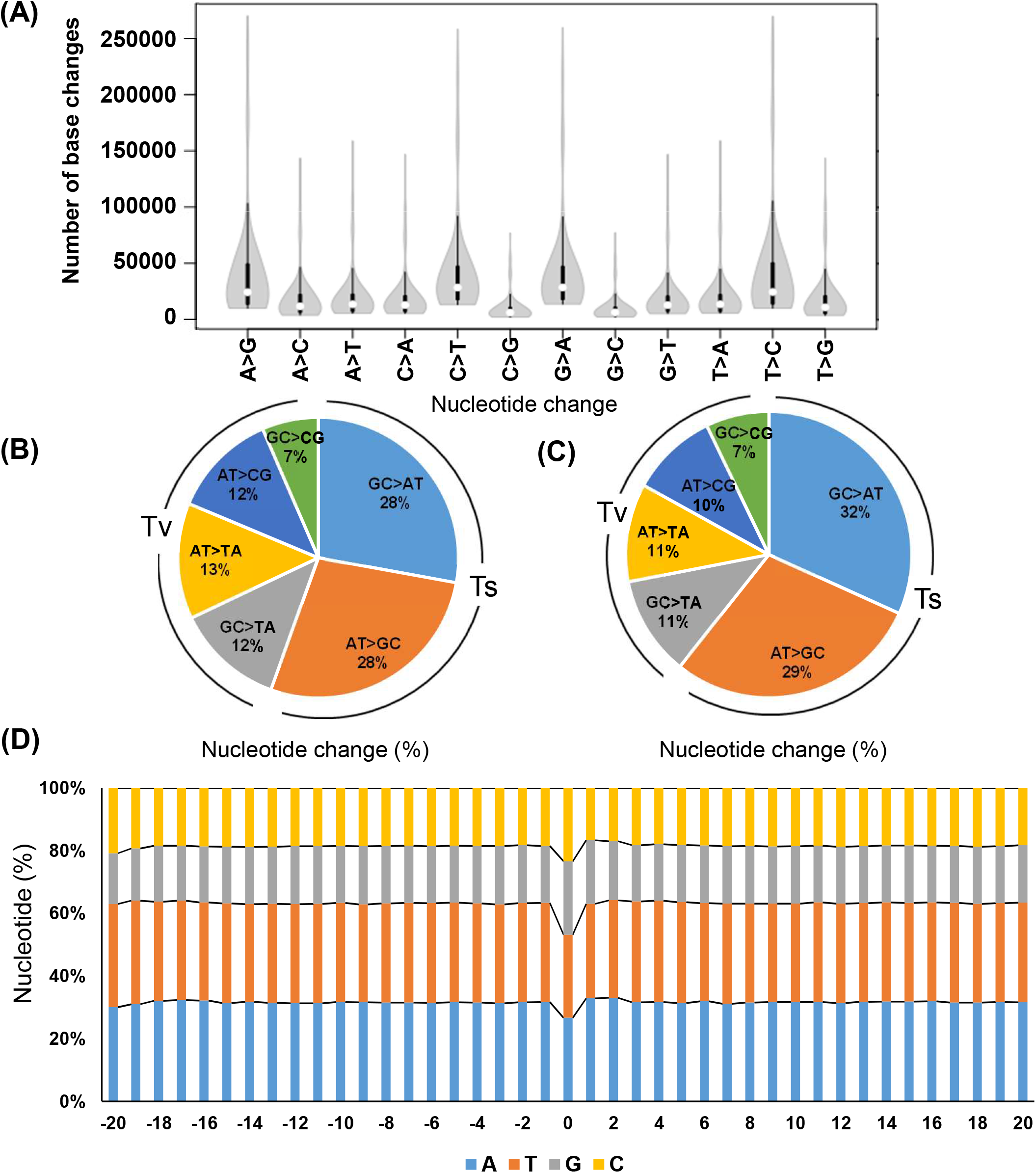
The spectrum of nucleotides changes in the mutagenized population. (**A**) The magnitude of changes in the different nucleotides in the mutagenized population. (**B-C**) The percent nucleotide changes in the whole genome (**B**) and CDS (**C**). (**D**) The flanking sequence (20 bp) on either side of the mutations in the genome. The 0 bp represents the site of mutation. The bars depict percent nucleotides at a given position in the sequence. The colors represent the individual nucleotides with annotations indicated below the graph. For details, see table S2

Purportedly, EMS-induced mutations are strongly biased to G/C-to-A/T transition (**Greene et al., 2003**), as EMS preferentially alkylates guanine to O^6^-ethylguanine, which mispair with T in place of C (**Ashburner, 1990**). To a reduced extent, EMS also mediates AT> GC transition by the alkylation at O^4^ of thymine (**Drake and Baltz, 1976**) (**Figure S1**). Therefore, we checked whether the digression to high A/T-to-G/C transition was restricted to a specific region or was omnipresent throughout the genome. The protein *C*o*D*ing *S*equences (CDS) analysis revealed a nearly similar mutation spectrum with 31.74% G/C-to-A/T and 28.93% A/T-to-G/C transitions. Correspondingly, the 7 to 11% transversion frequency in CDS was also closely similar to transversions in the whole genome (**Figure 1C**).

To ascertain if any bias exists in SNPs in the natural population, we chose the **Aflitos et al. (2014)** study for comparison because it contained SNPs from the present-day cultivars (53) and several wild relatives (31). Interestingly, we found a similar base change bias in the natural population, with 27.31% G/C-to-A/T and 26.88% A/T-to-G/C transitions **(Figure S2)**. The similarity between base changes frequency of EMS-mutant lines and tomato accessions shows that even for EMS, the retention of the mutations is akin to natural cultivars,

### Does EMS have any sequence bias?

We next examined whether EMS preferentially induces mutations at genic sites having any particular motif. We analyzed the nucleotide frequencies 20 bp upstream and downstream flanking to the 41 million identified SNPs. Remarkably we did not find any preferred genic motif or bases upstream or downstream of the mutated site. Nonetheless, our analysis showed that the mutated site had a high GC percentage (47% of GC-rich region) in comparison to the flanking sequences. (37% GC-rich region) (**Figure 1D**, **Table S2).**

### Nearly 31% of induced mutations in the population were unique

The penetrance of mutations showed a wide variation in the population. The mutation frequency ranged from 1/0.45 Kb to 1/11.05 Kb in mutant lines (**Table S3**). Allowing for the tomato genome size of 950 Mb, the average mutation frequency was 1/3.057 Kb, considering each line, on an average, harbored 311,101 SNPs. As EMS produces random mutations in the genome, we examined the distribution of SNPs in all 12 chromosomes of the population. Line-wise distribution of SNPs revealed that chromosome 11 had the highest SNPs in most lines. Oppositely, chromosomes 4 and 6 had the lowest SNPs (**Figure 2A**). However, the genome-wide SNPs distribution for all lines revealed that chromosomes 0 and 11 are densely populated with SNPs while chromosomes 3 and 8 are sparsely populated (**Table S4**). Both homozygous and heterozygous SNPs showed a random distribution across the genome. Foreseeably, the heterozygous SNPs were numerically higher than the other changes in the nucleotides (**Figure 2B**).

**Figure 2.**
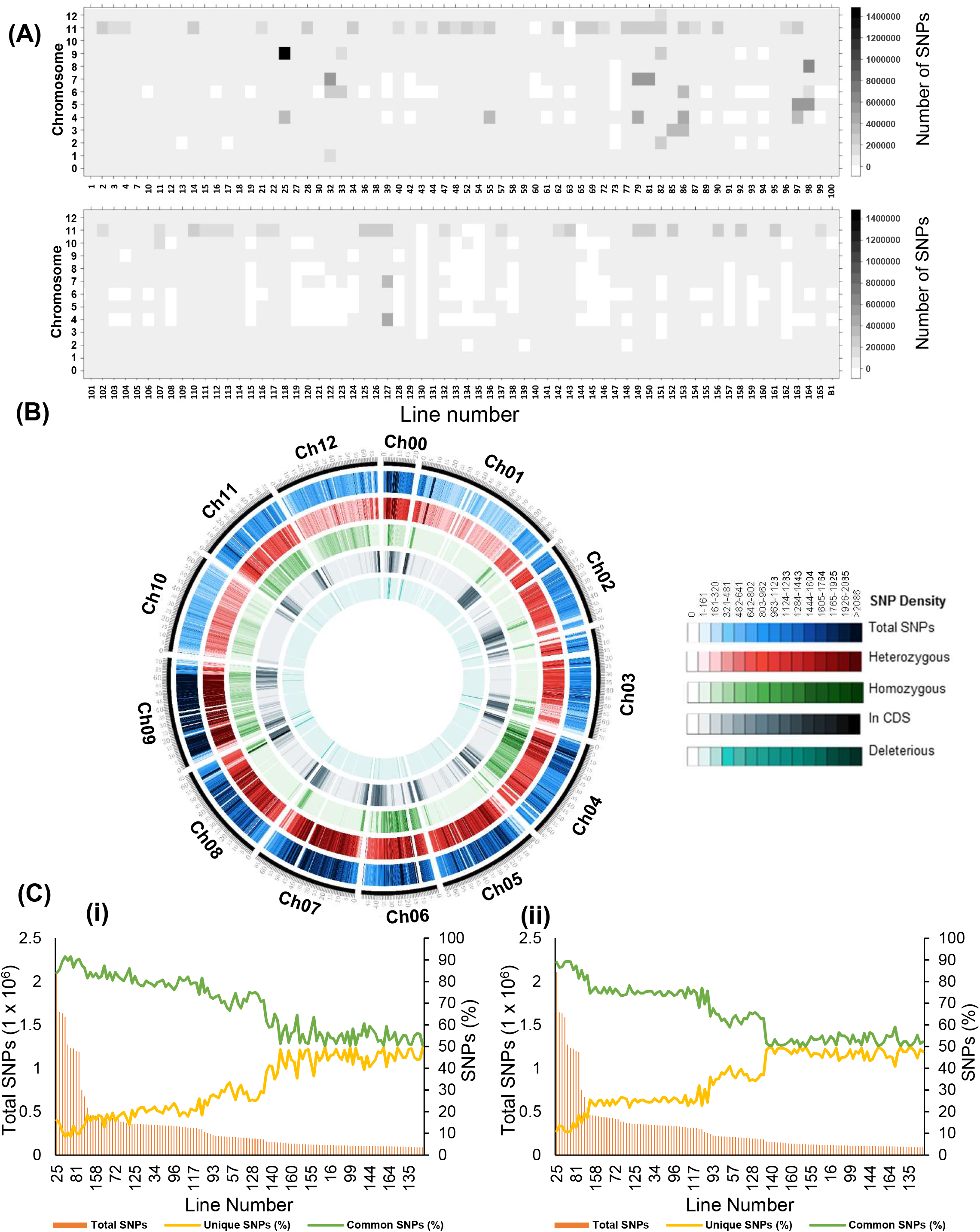
Distribution of the mutations across the genome. (**A**) Heatmap shows the distribution of the total number of SNPs in the 12 chromosomes in 132 mutant lines. (**B**) Circos plot showing the chromosome-wise distribution of different mutations in the mutant population. For details, see table S4. (**C**) The unique SNPs present in 132 mutant lines in comparison to 53 tomato cultivars (**i**) and 31 wild relatives of tomato (**ii**). Left axis-total SNPs, Right Axis-percent unique and common SNPs in mutant lines. Note percent unique SNPs are much lower in the mutant lines with the high density of the mutations. Lines are organized by decreasing number of total SNPs. For details, see table S5.

The exploitation of induced genic polymorphism of the mutant population greatly depends on the uniqueness of the altered SNP(s). We ascertained how many induced SNPs were novel than the genic polymorphism present in tomato cultivars. To do this, we compared the 41 million induced SNPs identified in our 132 mutant lines with 539 million naturally existing SNPs reported in 85 tomato lines (**Aflitos et al., 2014; Gupta et al., 2020a**). Moreover, in **Gupta et al. (2020a)** study, we realigned the sequences of 84 tomato lines to SL3.0 assembly, akin to this study. Compared to tomato cultivars and wild relatives, on average, 31.07% (*range* 8.53➔49.65%) and 35.37% (*range* 10.66➔49.96%) SNPs in 132-lines were unique in our lines, respectively (**Table S5**). Remarkably, the percent of unique SNPs in a mutagenized line is oppositely correlated with the density of mutations. The lines with lower mutation density had the highest percent of unique SNPs and vice versa (**Figure 2C, Figure S3). Prediction of effect of mutations on the encoded protein function**

To assess the impact of the individual mutation on the encoded protein function, the 41 million induced-SNPs were annotated using the SIFT4G ITAG3.2 genome reference database (**Vaser et al., 2016; Gupta et al., 2020a**). Based on the annotation, 91.5% SNPs were present in the intergenic region, 5.5% in the intronic region, 2% in the CDS region, and 1% in the UTR region (**Figure 3A**). Out of 2% SNPs in CDS, 36.3% led to synonymous (silent) mutations, 60.8% were nonsynonymous (missense) mutations, while 2.9% caused stop-gain/loss and start-loss, leading to truncation of the encoded protein (**Figure 3B**). On average, the CDS in a mutant line contained 2290 synonymous, 3834 nonsynonymous, 32 stop-loss, 136 stop-gain, and 16.5 start-loss mutations (**Table S6**). Besides, the mutant lines also had several deletions comprising both frameshift and non-frameshift InDels. On average, a mutant line harbored 694 frameshift insertions and 601 frameshift deletions (**Table S7**). We also found 454 unique mutations in 156 miRNAs, with heterozygous mutations (79%) constituting the majority (**Table S8).**

**Figure 3.**
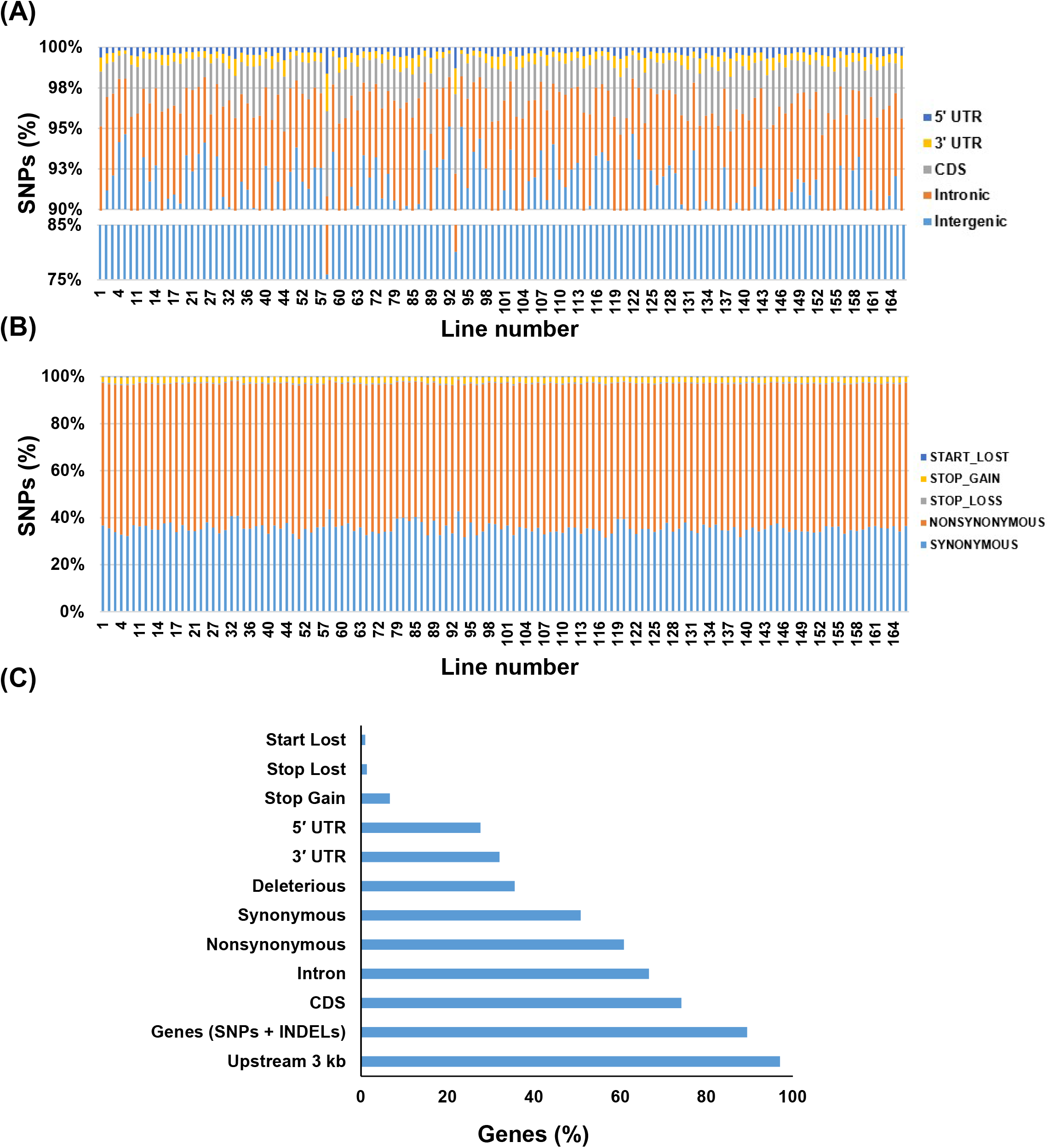
Distribution of mutations in the mutagenized population. (**A**) The relative distribution of mutations in different regions of the tomato genome in the population. For details, see table S3. (**B**) The distribution of different functional classes of mutations in the CDS region of the population. For details, see table S6. (**C**) The relative proportions of different functional categories of mutations, including mutations in promoters and introns.

In the 132 mutant lines, 89.4% genes harbored at least one SNP/InDel, with 74.2% having SNP/s in the CDS region. Among the SNPs in the CDS, 60.9% of genes harbored nonsynonymous mutations (**Figure 3A**). Our analysis pointed out that almost 97% of genes had at least one SNP/s in the 3 Kb upstream region of the gene (**Figure 3C**). Among these mutations, 35.6% of genes had mutations with SIFT score <0.05 and therefore predicted to be deleterious (**Ng and Henikoff, 2003; Kumar et al., 2009**). Consistent with the random nature, these deleterious mutations were distributed across the genome. A limitation of the SIFT is that it is geared to predict the effect of a single amino acid change on the protein function. The SIFT does not predict the influence of the start-loss, stop-gain, and stop-loss variants on the protein function. Indubitably, these variants can also be deleterious or affect the protein’s function. These variants were present in the range of 1% to 7% of the genes in the mutant population. The effect of the mutations in the intergenic region, particularly in the promoter, is not directly quantifiable like the ones affecting the amino acids (**Figure 3C**). Nonetheless, these mutations also are of great importance, as the variation in the Cis-regulatory region/promoter region can modulate gene expression and create trait diversity.

### Effect of mutations on individual codons and amino acids

The degeneracy of the genetic code protects the genome against the mutation loads caused by spontaneous mutations. Notwithstanding the genetic code degeneracy, nearly 62.8% of CDS mutations were nonsynonymous. Among the nonsynonymous changes, 4% belonged to the valine to isoleucine and vice-versa (V/I and I/V) and alanine to valine (A/V). Among the 37.2% synonymous mutations, the major changes were leucine-to-leucine (L/L) and serine-to-serine (S/S) (**Figure 4A**, **Table S9**). These two changes comprised around 9% of total amino acid substitutions. A high degree of synonymous changes in leucine and serine is expected, as these two amino acids are encoded by six codons.

**Figure 4.**
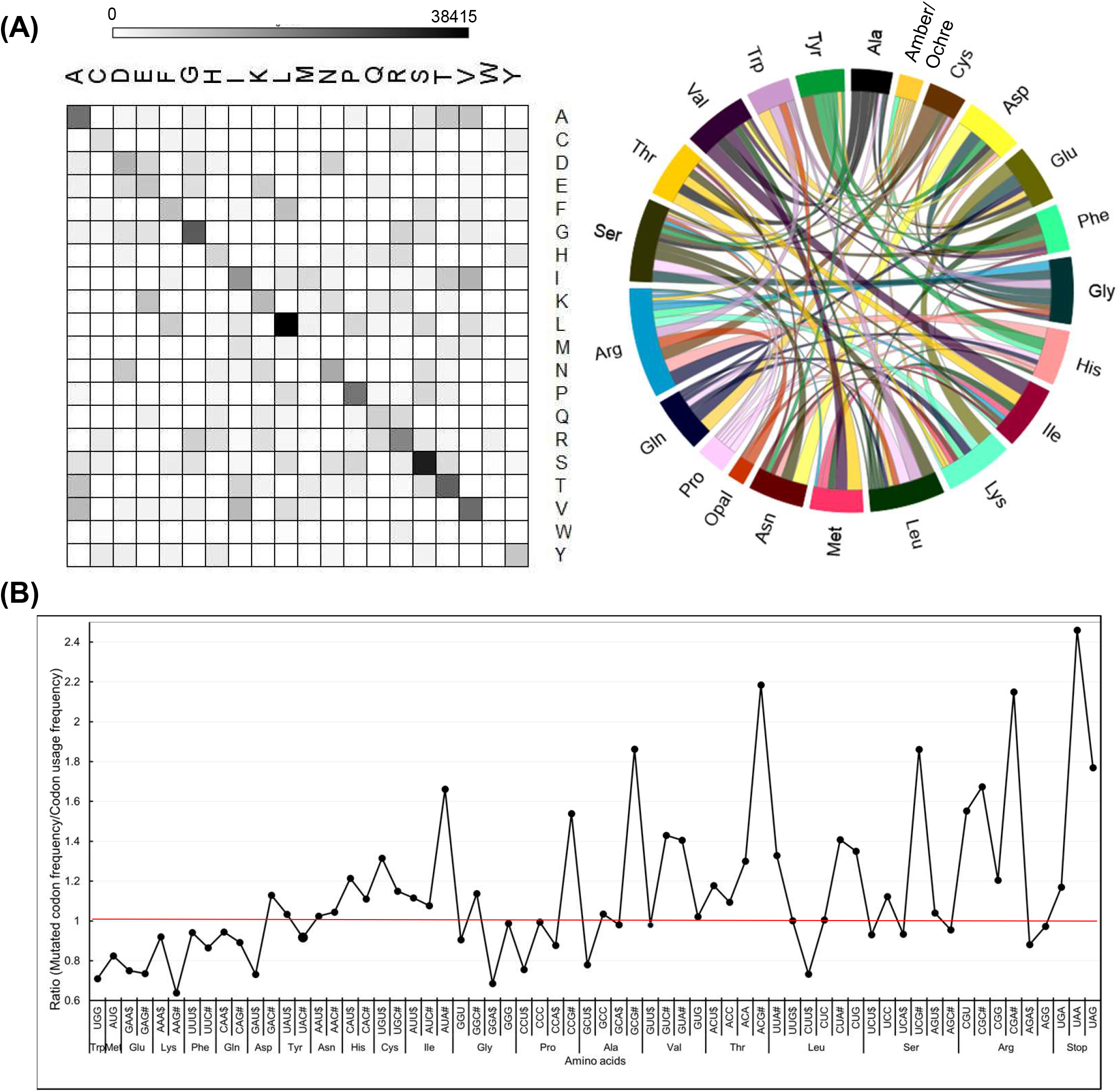
Frequency of mutations in different codons and amino acids. (**A**) Relative frequency of changes in individual amino acids in mutagenized population. All 168 possible amino acid changes were observed with varying frequency. For details, see tables S9 and S12. (**B)** The ratio between the frequency of a mutated codon in the mutagenized population and its normal frequency in tomato. The most preferred codon as per the tomato codon usage table (https://solgenomics.net/documents/misc/codon_usage/codon_usage_data/l_esculentum_codon_usage_table.txt) is marked with the $ dollar sign, and the least preferred codon is marked as # hashtag after the codon letter in the graph. The individual ratios are given in Table S10.

We compared whether the mutagenicity of codons had any relationship to their usage in a protein by comparing with codon usage frequency in tomato. We checked it by calculating the ratio between the frequency of a mutated codon and its usage in wild-type tomato. We considered the most used codon for an amino acid as the preferred codon. Interestingly, for amino acids having higher degeneracy of 3-4 codons, the preferred codons are the least prone to mutagenesis (ratio ≤1). Conversely, the least preferred codons are most prone to mutagenesis (ratio ≥1). Interestingly, even for the stop codons, the ratio for the most preferred codon, UGG, was lower (1.04), while less preferred codons (UAA-1.86, UAG-1.82) had a higher ratio. (**Figure 4B, Table S10**). Remarkably, the analysis of tomato cultivars also shows that the codon least preferred is the most prone to mutation, with a near overlap in the pattern of mutagenized lines and cultivars (**Figure S4,** Table S11).

Markedly, the amino acids encoded by two codons do not show the above mutagenicity bias as the ratios are nearly similar and largely remain ≤1. Comparing the ratio of the frequency of mutated codons with that of codon usage revealed that methionine (0.73) and tryptophan (0.77) are the least prone to mutagenesis. Since the methionine and tryptophan lack codon degeneracy, any mutation leads to a nonsynonymous change.

The dominance of transitions over transversions in the overall mutational spectrum is strongly seen in the CDS mutations. The theoretically predicted ratio of mutations for an amino acid calculated with the assumption that mutations are random strongly deviates from the observed mutations (**Figure 4C**, **Table S12**). For illustration, the mutation from valine to alanine, where the middle codon changes from T to C, signifying transition, has a ratio > 1 (24.49/16.66= 1.46). In contrast, the mutation from valine to glycine, where the middle codon changes from T to G, signifying transversion, has a ratio < 1 (8.39/16.66= 0.50). The above pattern of difference between theoretical and actual distributions of transitions or transversions is consistently observed for most amino acids (**Table S12**). The average ratio of transitions (1.80) was nearly 2.38 fold higher than that of transversions (0.756).

### Metabolic pathways affected by mutations

Gene Ontology (GO) analysis revealed a broad range of the categories bore nonsynonymous and synonymous mutations, including mutations in the UTR. (**Figure 5A, Figure S5, Table S13**). In the mutant population, nearly 40% genes bore no nonsynonymous mutations (**Figure 3C**). Across the different GO categories, the absence of mutations was mainly in housekeeping genes related to the gamut of the essential cellular processes. For instance, no mutations were detected in GO categories such as tetrahydrobiopterin synthesis, t-RNA 3′ end processing, mitochondrial respiratory chain complex, and ATPase activity regulation.

**Figure 5.**
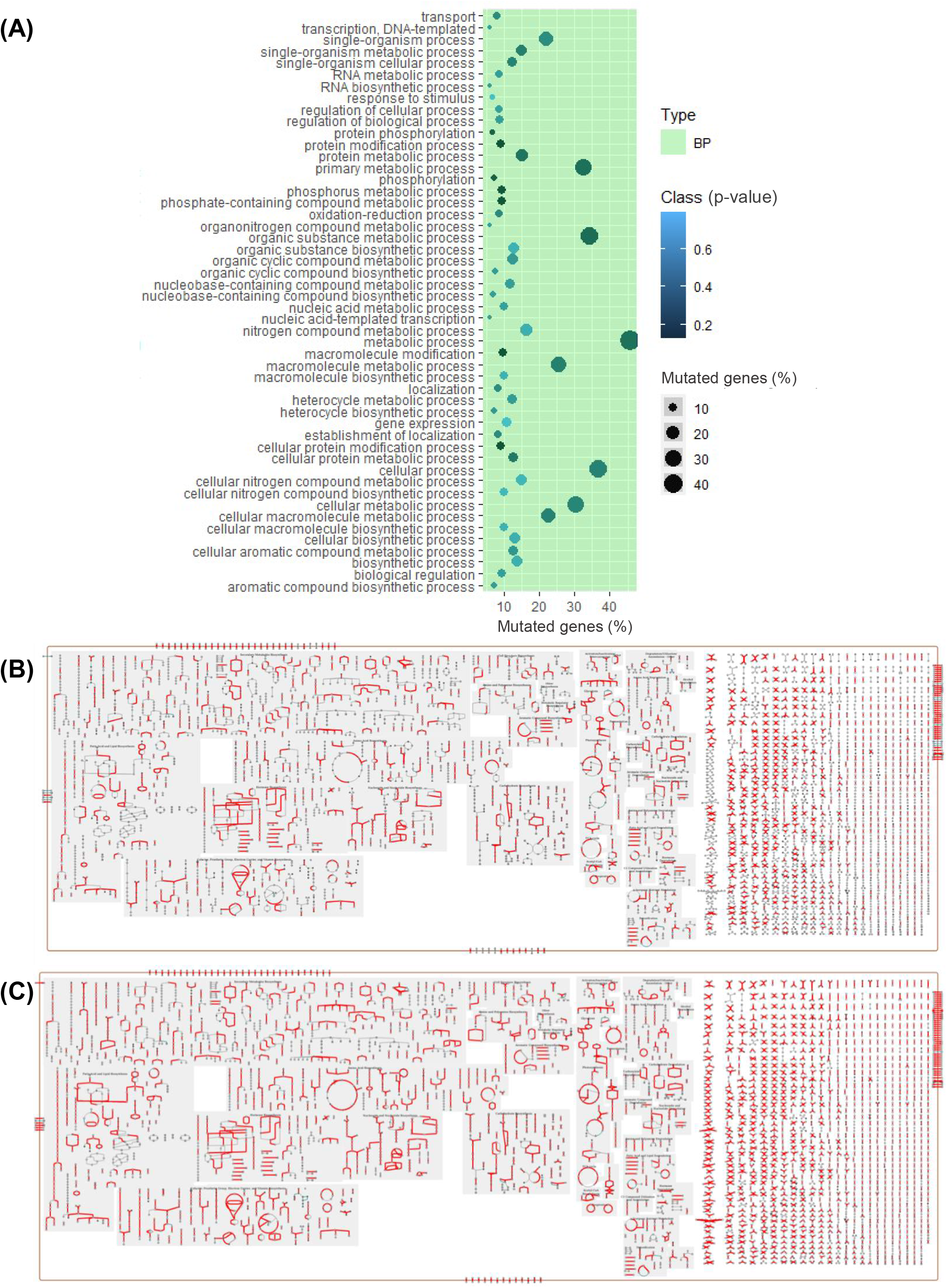
Distribution of mutations in different GO categories and on the metabolic pathway. (**A**) Top 50 GO categories (Biological process-BP) with the high frequency of mutations. Note, in none of the GO categories; the percent mutations exceed 50%. For details, see Table S13. (**B**) Tomato metabolic pathway steps marked with deleterious mutations present in the mutagenized population. For details, see Table S15. (**C**) Tomato metabolic pathway steps marked with nonsynonymous mutations present in the mutagenized population. Gene Id’s were mapped to the tomato metabolic network from Plant Metabolic Network Database (https://pmn.plantcyc.org/overviewsWeb/celOv.shtml?orgid=TOMATO).

To specifically evaluate the influence of mutations on a wider scale, we mapped the genes predicted by SIFT to have loss-of-function (LOF) on the tomato metabolic pathway. Out of the 7991 genes currently assigned in LycoCyc to the tomato metabolic pathway (https://solcyc.solgenomics.net/organism-summary?object=LYCO, ver. 3.8. ITAG3.2), 2861 genes were mapped with LOF (**Figure 5B**). The mapping of all nonsynonymous mutations on tomato metabolic pathways highlighted that most pathways had one or more mutant genes that may affect their operation (**Figure 5C**).

We then specifically examined the distribution of mutated genes on three important metabolic pathways, viz. tetrahydrofolate biosynthesis, carotenoid biosynthesis, and plant photoreceptors/circadian regulation. We found 3 deleterious mutations affecting the tetrahydrofolate pathway **(Figure S6**), 15 deleterious mutations in the light-signaling pathway (**Figure S7**), and 11 deleterious mutations in the carotenoid biosynthetic pathway (**Figure 6A**). To validate whether a SIFT-predicted deleterious mutation alters the phenotype, we selected two mutants, one from the carotenoid pathway and one from the photoreceptors.

**Figure 6.**
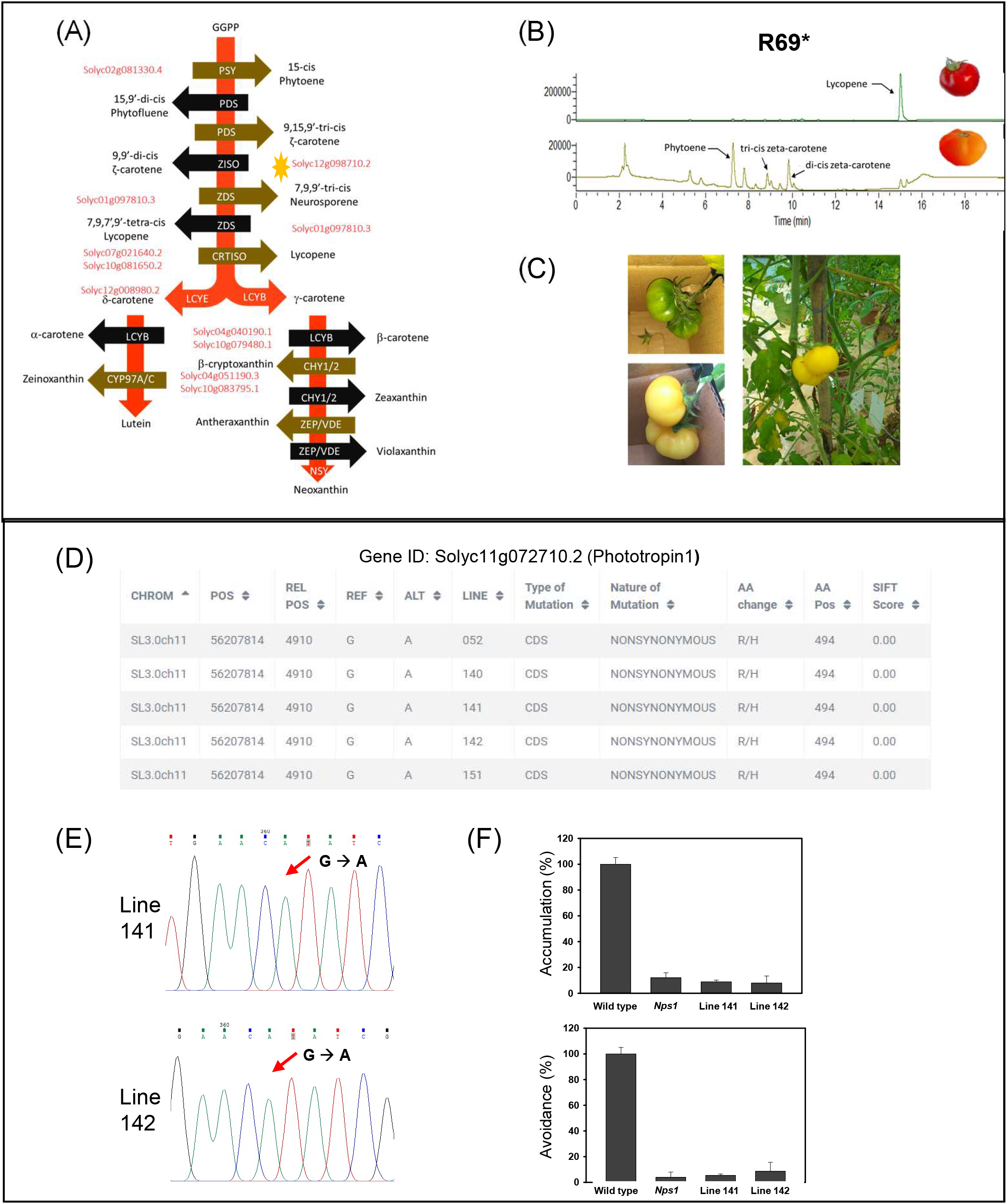
Validation of light signaling and carotenoid biosynthesis mutants. (**A**) Carotenoid biosynthesis pathway in tomato showing the 11 deleterious mutations present in the population (Gene Ids are in red, *ziso* mutation is marked with *). (**B**) The chromatograms show the loss of lycopene formation in the *ziso* mutant (R69*). Note, unlike wild type, the mutant forms little lycopene but shows tri-ci-ζ-carotene and di-cis-ζ-carotene (bottom panel), which are present in traces in wild type (top panel). (**C)** The loss of ζ-carotene accumulation in *ziso* fruit shielded from light. The fruit was shielded from light at the mature green stage, and photographs were taken after 12 days. The absence of light massively reduced the formation of tri-ci-ζ-carotene and di-cis-ζ-carotene in fruits. (**D**) Screenshot from ITGV database, showing the lines containing R494H substitution in the *phototropin1* gene. **Note:** In recent ITAG3.2 annotation, the *Nps1* mutation is located at R494H. (**E**) The homozygosity of mutations in lines 141 and 142 was validated by Sanger sequencing. (**F**) The chloroplast relocation response in leaves of mutant lines 141 and 142. Both lines show near-total loss of chloroplast accumulation and avoidance response, similar to *Nps1* mutant. (Abbreviations: PSY, phytoene synthase; PDS, phytoene desaturase; ZISO, ζ-carotene isomerase; ZDS, ζ-carotene desaturase; CRTISO, carotenoid isomerase; LCYE, lycopene ε-cyclase; LCYB, lycopene β-cyclase; CYP97A/C, cytochrome P450 monooxygenase; CHY1/2, beta-carotene hydroxylase; ZEP, zeaxanthin epoxidase, VDE, violaxanthin epoxidase; NSY, neoxanthin synthase)

### Characterization of a **ζ**-carotene isomerase mutant

The carotenoids act as photosynthesis accessory pigments and protect the photosynthesis machinery from photooxidation. Therefore, the mutants compromised in the early steps of carotenoids biosynthesis are lethal, with the notable exception of carotenoids isomerization genes such as ζ-carotene isomerase (*ZISO*) and carotene isomerase (*CRTISO*). The carotenoid isomerization catalyzed by ZISO and CRTISO can also be catalyzed by light. Thereby photosynthetic activity in green tissues is not comprised in these mutants. However, the light fails to penetrate the deeper tissue layers of tomato fruits. Therefore, carotenoid biosynthesis gets stalled at the conversion of tri-cis-ζ-carotene to di-cis-ζ-carotene in ZISO mutant resulting in orangish-red fruits.

We identified a deleterious homozygous mutation in the *ZISO* gene in the B1 line, harboring a stop codon at the 69^th^ amino acid position. All progeny plants from the B1 line bore orangish-red fruits, a characteristic shared with reported *ziso* mutants of tomato (**Fantini et al., 2013**). Consistent with the *ziso* mutant being compromised in carotenoids isomerization, the fruits had higher amounts of tri-ci-ζ-carotene and di-cis-ζ-carotene, confirming that the mutant was indeed comprised in ZISO activity. The tri-ci-ζ-carotene and di-cis-ζ-carotene do not accumulate in wild-type fruits, as light along with ZISO and CRTISO enzymes convert them to downstream carotenoids, mainly lycopene (**Figure 6B**). To reduce light-mediated isomerization of carotenoids, we covered on-vine the mature-green *ziso* mutant fruits with black sheets. The enclosed fruits were yellow-colored and had reduced levels of tri-ci-ζ-carotene and di-cis-ζ-carotene, and low lycopene, confirming that the covering of fruits substantially blocked light-mediated carotenoid isomerization (**Figure 6C**).

### Characterization of a phototropin1 mutant

In tomato, the mutation in the phototropin1 gene changing arginine to histidine at 495^th^ amino acid (Arg495His, now revised to R494H as per ITAG 3.2) dominantly blocks the light-induced chloroplasts accumulation and avoidance response in leaves (**Sharma et al., 2014**). We found the same genic variant in five mutant lines viz. 052, 140, 141, 142, and 151. Two mutant lines, 141 and 142, had the mutation in the homozygous state. We examined the chloroplast relocation responses in homozygous 141 and 142 lines, wild type (Arka Vikas), and *Nps1* mutant. Consistent with the dominant-negative effect of Arg494His mutation, the chloroplast relocation response in 141 and 142 lines was blocked similarly to the *Nps1* mutant and its backcrossed progeny. (**Figure 6D-F, Figure S8**).

### Web-searchable access to mutations

To make this comprehensive mutant resource and its corresponding data available to the public, we made an open-access database called ITGV (Induced Tomato Genomic Variations) database (http://psd.uohyd.ac.in/itgv/). The users can search the ITGV database by gene id/name or mutant line. The search page provides the results with the promoter and gene region mutations. For the gene-specific mutations, SIFT annotation is also provided along with the SIFT score (**Figure 7**). Users can also visualize the SNPs and InDels through the genome browser “Jbrowse.” Mutation information of all the lines can also be downloaded from the ITGV. The users can request mutant seeds, the details of which are provided on the ITGV website.

**Figure 7.**
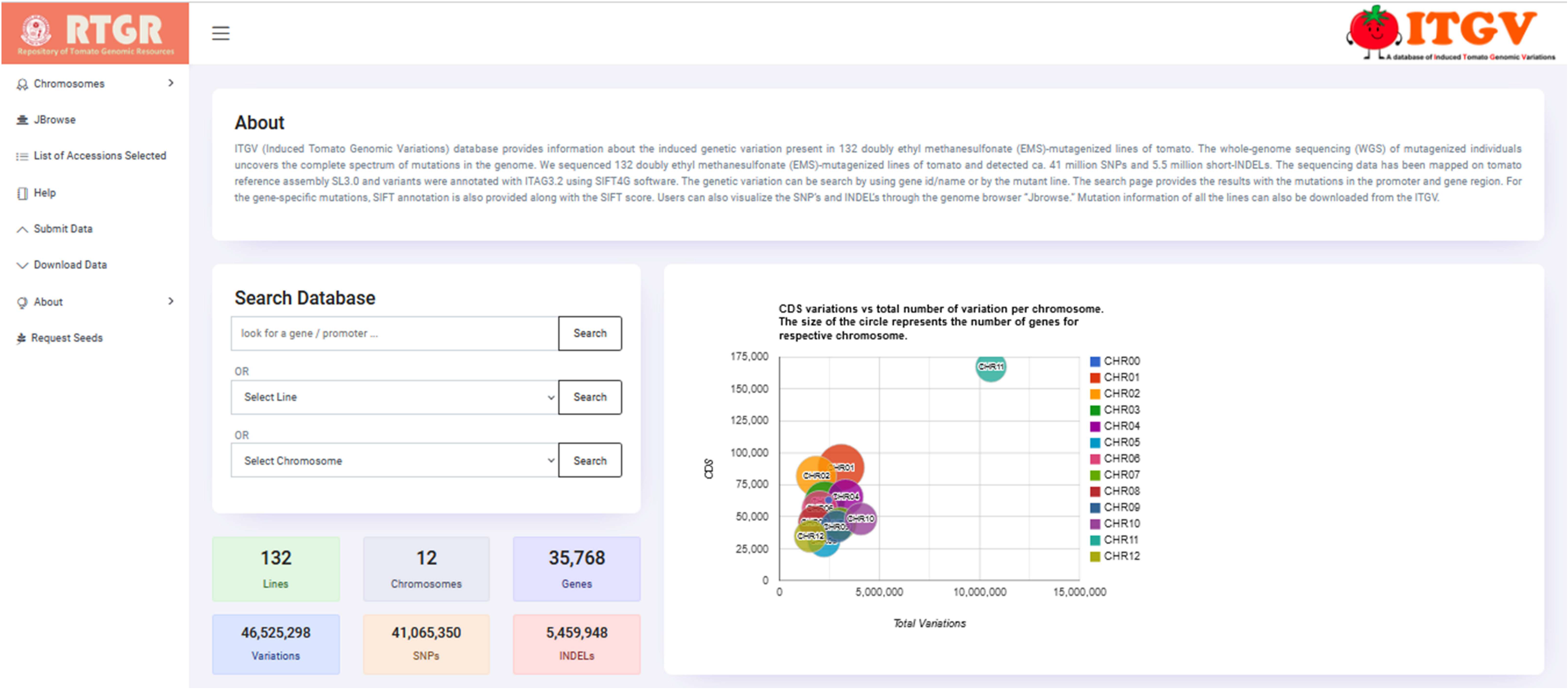
Screenshot of the tomato mutant database for variants visualization, variant information, and annotation (http://psd.uohyd.ac.in/itgv/)

## Discussion

### WGS uncovered a very high mutation density in tomato

In this study, we sequenced and cataloged an EMS-mutagenized tomato mutant population. The genome-wide analysis of mutations brought forth an unexpected aspect discordant with earlier reports. The observed average mutation density of 1/3.05 Kb is many-fold higher than earlier reports. Overall the magnitude of mutations was >100,000 per mutant line. The high mutation density also entails that a relatively small population is needed to mutate all the genes in tomato. The analysis of tomato EMS-mutagenized TILLING mutants revealed a mutation density range from 1/322 to 1/1710 Kb (**Gady et al., 2009; Minoia et al., 2010; Piron et al., 2010; Okabe et al., 2011, 2013; Okabe and Ariizumi, 2016**). In an earlier TILLING study, we found a mutation density of 1/367 Kb in the Arka Vikas cultivar (**Gupta et al., 2017**). Ostensibly, the WGS being more robust uncovered the real quantum of mutations in our population. The higher mutation density in tomato is also contrary to the notion that only polyploid species such as wheat (**Uauy et al., 2009**) or polyploid Arabidopsis (**Tsai et al., 2013**) can tolerate the mutation load owing to extra copies of the genome.

It remains possible that in conventional TILLING, due to the widely varying efficiency of CEL-I to cleave different mismatches (C/C ≥ C/A ∼ C/T, G/G > A/C ∼ A/A ∼ T/C > T/G ∼ G/T ∼ G/A ∼ A/G > T/T; **Oleykowski et al., 1998**), the bulk of mutations escape detection. Similarly, the mutations can escape detection in NGS-based TILLING due to the inadequacy of available software(s) to identify precisely mutations in 64X DNA pooling (**Gupta et al., 2017**). In whole-genome sequencing (WGS) using individual mutant lines, the variant detection in a mutagenized population is more robust, particularly with sequencing coverage higher than 15X (**Thompson et al., 2013; Krasileva et al., 2017**). The higher sequencing depth of 30X allowed us to detect a larger number of variants ranging from 1/0.45 Kb to 1/11 Kb in mutant lines.

In maize using whole-exome sequencing (WES) at 14.82X coverage for exons (Intron 9.48X, Promoter 7.35X coverage), mutation density was 1/48 Kb (**Lu et al., 2018**). Considering that WES was carried out in M_1_ maize plants obtained from mutagenized pollens, the mutation frequency (1/48 Kb) was practically half, as only a single haploid genome was mutagenized. In Cadenza wheat M_2_ population, WES at 29.01X coverage revealed the average mutation frequency of 1/30 Kb (**Krasileva et al., 2017**). In maize, like humans (**Belkadi et al., 2015**), a comparison between WES and WGS revealed that WGS is more robust in mutation detection than WES, even in the exome. Compared to the WES of the Bp001 mutant maize line, the WGS detected nearly 2-fold higher mutations in the same exonic region (**Lu et al., 2018**). Thus, it can be inferred that usage of WGS with ≥30X coverage allowed the detection of a far higher mutation density than other studies.

Interestingly, the lines bearing high mutation density also share a high percent of SNPs present in 53 tomato cultivars. It can be construed that the SNPs present in tomato cultivars arose by spontaneous mutations and were not subjected to purifying selection. In all likelihood, the EMS mutations in these SNPs had a higher probability of being retained and passed to the next generation. Considering that unique SNPs at maximum were <50% underscores the importance of purifying selection.

### EMS also caused a high frequency of AT> GC transitions

Several studies using WES or WGS restricted mutation analysis to GC>AT transitions (**Lu et al., 2018; Addo-Quaye et al., 2017**), though SNPs other than GC>AT contributed to 51% and 54.9% of total SNPs in Sorghum (**Addo-Quaye et al., 2017**) and maize (**Lu et al., 2018**), respectively. In this study, WGS revealed that GC>AT transitions comprised only 30% of overall mutations in the mutant population. In an earlier study, the amplicon sequencing of the progenitor of our mutant population revealed 65% GC>AT transitions (**Gupta et al., 2017**). Considering that we used a doubly mutagenized EMS [120 mM (1.5%); 2X) population, the remutagenesis likely reduced the frequency of GC>AT transitions. Seemingly higher dosage of EMS reduces GC>AT transition in tomato, as reported by **Minoia et al. (2010),** whereupon increase from 0.7% to 1% EMS reduced GC>AT transition from 60% to 28%. The mutations in our population showed a substantial overlap with SNPs present in the tomato cultivars. Since the SNPs have no nucleotide bias, the overlap emphasizes that non GC>AT transitions in the mutant population resulted from EMS-treatment.

The selection of GC>AT transitions as the genuine mutations are based on the premise that EMS introduces an alkyl group at O^6^-guanine, which leads to mispairing of G to T during DNA replication. It is believed that, unlike humans, O^6^-alkylguanine is not repaired in plants, as the plants reportedly lack O^6^-alkylguanine-DNA-alkyltransferase activity (**Leitao, 2011; Pegg, 2011**). It remains to be determined whether the reported removal of O^6^-alkylguanine in *Vicia faba* root tips (**Baranczewski et al., 1997a,b**) uses an alternate mechanism such as base excision repair (**Manova and Gruszka, 2015**).

The EMS-driven non GC>AT transitions are considered the side effect of mutagenesis and often are not included as potential mutations (**Addo-Quaye et al., 2017; Lu et al., 2018**). It may be imprudent to ignore these mutations, as WGS does not have a nucleotide bias, and at high coverage such as 30X used in our study, the error rate is extremely low. EMS-treatment can induce secondary mutations due to errors in DNA repairs, including DNA breakage, as evident by the presence of InDels in the mutant population. In crop plants, the frequency of mutations other than GC>AT widely vary with 10% in Arabidopsis (**Greene et al., 2003**), 30% in rice (**Till et al., 2007; Henry et al., 2014**), and 54.9% in maize (**Lu et al., 2018**). The WGS of EMS-mutagenized Toxoplasma revealed that ∼74% of mutations were in the A/T base pair (**Farrell et al., 2014**). The WGS of EMS-treated MicroTom lines revealed 39%-76% GC>AT transition (**Shirasawa et al., 2016**), while the WES of 95 tomato mutants displayed only 20.7% GC>AT transitions (**Yano et al., 2019**). We guess that the wide difference in GC>AT transition is a species-specific phenomenon perhaps related to the difference in the repair efficiency.

### O^4^-alkyl-thymine may be the causative agent for AT> GC transitions

The AT> GC transition arises from EMS-mediated the alkylation at O^4^ of thymine (**Drake and Baltz, 1976**), which can mispair during DNA replication leading to mutagenicity. In conformity with the above, in *E. coli* and human cell lines, the incorporation of O^4^-alkyl-thymine in DNA results in a large number of T➔C mutations (**Wang et al., 2015; Wu et al., 2016**). It is plausible that in plants, the formation of O^4^-alkyl-thymine may lead to T➔C mutations during DNA replication.

Contextually, it is likely that the higher degree of A/T-to-G/C transitions observed in this study reflects the mutagenicity of O^4^-alkyl-thymine residues. In this and an earlier study (**Gupta et al., 2017**), the Sanger sequencing validated both GC>AT and AT>GC transitions with nearly the same frequency. Thus it can be surmised that in parallel to GC>AT transitions, EMS also induces the AT> GC transition, albeit at a lower frequency. Compared to EMS-mediated O^6^-alkyl-guanine formation, the formation of O^4^-alkyl-thymine is much less (**Leitao, 2011**). The lower frequency of AT> GC transition is consistent with the said EMS efficiency and augurs with the increase in AT> GC transitions with higher EMS dosage to tomato (**Minoia et al., 2010**).

### The least preferred synonymous codons are most prone to mutagenesis

The degeneracy of genetic code predisposes that the synonymous codons for an amino acid vary widely in frequency (**Ikemura, 1985**). It is believed that among the highly degenerate codons, some codons are preferred over others because they are translated more efficiently and accurately (**Hershberg and Petrov, 2008, 2009**). Another view is that evolutionary selection favors preferred codons over other minor codons, while mutational pressure and genetic drift allow the minor codons to persist (**Bulmer, 1991**). Contrarily, in moss, it was suggested that weak natural selection for translational efficiency shapes the codon bias rather than mutational bias (**Stenøien, 2005**). Our results bring a different paradigm to the codon usage bias. It can be construed that the preferred codons are least mutagenic to EMS, as these are the main codons for translating critical proteins such as ribosomal proteins, elongation factors, and t-RNA. Conversely, the least preferred codon has the highest propensity to mutagenesis. Seemingly, mutation bias plays a role in selecting the preferred codon in tomato. The codon most preferred in natural selection is strongly disfavored for mutations.

The classification of mutations across the genetic code table reveals that the transversions generate more nonsynonymous mutations than the transitions. Yet, transitions have a ratio higher than the theoretically possible ratio at mutated amino acid levels, while transversions show a converse pattern. Considering that the nonsynonymous transitions are purported to be less deleterious (**Zhang, 2000; Lyons and Lauring, 2017**), the transitions have more likelihood of being retained in progeny. The less deleterious effect of transitions could be related to their influence on protein function, as transitions do not cause drastic changes in amino acid physicochemical properties such as polarity, charge, and size (**Zhang, 2000**).

### Housekeeping genes are recalcitrant to EMS-mutagenesis

In mammals, approximately 35% of genes are essential for survival and housekeeping (**Dickinson et al., 2016**), and the same is likely for plants. In *Plasmodium falciparum,* large-scale insertional mutagenesis revealed that nearly 50% of genes are essential for optimal growth (**Zhang et al., 2018**). Since many GO categories showed no nonsynonymous mutations, these genes are presumably essential for tomato. Our results are in conformity with large-scale mutagenesis in *Caenorhabditis elegans*, where too the essential genes lacked mutated alleles (**Thompson et al., 2013**). The lack of mutations in housekeeping or essential genes in tomato indicated that these genes are largely recalcitrant to mutagenesis. It can be construed that, similar to spontaneous mutations in nature (**Monroe et al., 2022**), the EMS-induced mutations in essential genes are subjected to strong purifying selection and eliminated. It remains to be determined whether the epigenomic state of essential genes, as observed in Arabidopsis, reduces the rate of mutations (**Monroe et al., 2022**). It may be noted that our GO analysis represents only the coding mutations. Nonetheless, the mutations in the noncoding region may also influence the function and/or expression of essential housekeeping genes.

### Nearly 3000 deleterious mutants mapped on different metabolic pathways

The repertoire of mutations identified in our study is valuable to identify the function of unassigned as well as known genes. In particular, the mutations in genes modulating metabolic pathways can be potentially used either singly or in combination to examine the influence on metabolome/proteome and plant phenotype. In maize (**Lu et al., 2018**) and sorghum (**Addo-Quaye et al., 2017**), mutations in gibberellin (GA) biosynthesis were preferentially selected to validate the mutations identified by WES and WGS, respectively. The above mutants compromised in the GA biosynthesis pathway were dwarf due to reduction in GA levels; thereby, the phenotype was rescued by GA. We selected two highly penetrant mutants, the ZISO enzyme that executes carotenoids isomerization, a key step in carotenoids biosynthesis, and phototropin1, which mediates chloroplast accumulation to optimize photosynthesis under weak light.

Though a single copy gene encodes ZISO, the knockout mutation is not lethal, as the carotenoid isomerization can also be photochemically carried out by light. Since insufficient light penetrates the deeper layers of fruits, *ziso* mutant accumulates di- and tri-cis-ζ-carotene, while in wild-type, these are below the detection limits. Consequently, the lycopene level in the *ziso* fruits is considerably reduced than wild type. Further, in covered *ziso* fruits, lycopene accumulation is massively reduced due to blockage in light, which supports the role of ZISO in carotenoid isomerization.

The dominant-negative *Nps1* mutation (Arg494His) in phototropin1 blocks chloroplasts’ relocation responses in the mutant leaves. The chloroplasts stay at the bottom of mesophyll cells and do not move toward weak light or move away from strong light (**Sharma et al., 2014**). Like the *Nps1* mutant, two independent homozygous phototropin1 mutants (Arg494His) lines lacked the chloroplast relocation response. The loss of chloroplast accumulation in BC_1_F_2_ lines further supports that Arg494His mutation has similar dominant-negative action in Arka Vikas as reported for *Nps1* in Ailsa Craig background.

Screening our mutant resource using new and uncharacterized genes may uncover the phenotypes that were not covered by the forward genetics, particularly those leading to metabolic changes but may not have a phenotype. To that effect, we provide a cellular overview of the metabolic pathway where mutations may be affecting steps of a given pathway. In addition, the mutant resource provides scope for silencing a particular pathway by combining the mutations from different lines into the wild-type background.

### WGS provides a broader repertoire of mutants than WES

Unlike Arabidopsis, the function of the majority of genes in tomato remains unexplored due to the lack of mutants and t-DNA/transposon tagged lines. The availability of a gene-indexed mutation database bridges this gap for tomato functional genomic analysis (http://psd.uohyd.ac.in/tgv/). The high density of mutations in our population, in essence, provides multiple alleles for several genes. These allelic variants can be used for analyzing the function of a selected gene or a group of genes. The potential effect of mutations on the protein function can be assessed using the SIFT prediction incorporated in the database. Since SIFT presages the extent of the deleterious effect of the mutation, the influence of stronger alleles on a phenotype/response can be compared in tandem with weaker alleles. While SIFT predicts the loss-of-function mutations, lamentably similar tools for predicting the gain-of-function mutations are amiss.

Though the genic mutations are the main contributors to the phenotypes, emerging evidences indicate that promoter, UTRs, and intronic mutations also affect phenotypes by influencing gene expression. Likewise, mutations in miRNA genes affect the post-transcriptional regulation of several genes. Even the synonymous mutations are worth to be examined, as these mutations often influence a trait due to the organismal bias for codon usage. It is reported that synonymous mutations constitute ca. 1/3^rd^ of CDS mutation (**Henry et al., 2014; Thompson et al., 2013**). Our analysis is also consistent with this, as 37.39% of mutations in CDS were synonymous.

Unlike SIFT, the influence of the above mutations cannot be a priori predicted; the confirmation of their mutagenicity requires a detailed phenotype/biochemical examination. Anyway, the WGS used by us is superior to the WES, as it brings out the variants that WES does not discover. The WES largely excludes intronic variants, and promoter mutations, as the emphasis is focused on the CDS region. It is now increasingly becoming evident that alteration of phenotypes is not restricted to genic variants, but intergenic regions variants too contribute a lot.

### WGS provides a broader resource for trait improvement

Functional genomic analysis of noncoding mutations has emphasized their role in key biological processes in plants (**Li et al., 2017**). In tomato, the noncoding RNAs, circular RNA, and miRNA regulate protein-coding gene expression through diverse mechanisms (**Ma et al., 2020; Zuo et al., 2020**). Our WGS analysis revealed a significant number of mutations in noncoding regulatory elements encompassing promoters, introns, 5′- and 3′-UTRs. Our resource includes mutations in miRNA, and several of these reportedly regulate development in tomato (**Zuo et al., 2020; Ma et al., 2020**). The availability of noncoding mutations expands the mutation spectrum to discover elements regulating gene function or a biological pathway in tomato.

One may argue that owing to high background mutations, the assessments of gene function may be difficult. However, for genic mutations, the SIFT predictions are quite reliable and can be used as starting point. Additionally, examination of multiple alleles of a given gene also generally overshadows the effect of the background mutations. The mutations can be first validated by Sanger Sequencing. To confirm genotype and phenotype cosegregation, the Mutmap approach can be used (**Garcia et al. 2016**), followed by backcrossing During backcrossing at the seedling stage, the mutated gene can be identified in the heterozygous and homozygous state using CEL-I, a mismatch-specific endonuclease. The heterozygous plants carrying the mutation can be recurrently backcrossed with the desired cultivar (**Sharma et al., 2021**; F**igure S9**). With the mutated gene itself being a marker, in BC_4_F_1_ generation, 98% of mutations are eliminated (**Hospital, 2003**). In BC_4_F_2_, the homozygous mutant plants can be subjected to WGS to select the nearest isogenic BC_4_F_2_ line to the foreground cultivar.

### Induced mutagenesis by EMS vis-à-vis Genome editing

Recently the CRISPR/Cas9-based genome-wide editing for CDS has been applied to rice (**Lu et al., 2017; Meng et al., 2017**) and soybean (**Bai et al., 2020**). The genome-edited mutagenesis requires plant transformation with a large number of gRNA constructs, designed *a priori* to disrupt the function of selected genes. The constructs are individually made and multiplexed-pooled to ease the large-scale transformation. The genome-edited plants are identified using standard protocols of identification of transgenic plants and validation of editing.

In rice (**Lu et al., 2017; Meng et al., 2017**) and soybean (**Bai et al., 2020**), mutagenesis by genome-editing mainly generated deletions. A more rigorous analysis in maize revealed that most edited genes had deletion (60%) than insertion (32.5%). In the remaining 8% of genes, the 2/3^rd^ had transversions, and 1/3^rd^ had transitions (**Liu et al., 2020**). Broadly, genome-edited mutagenesis is akin to fast-neutron mutagenesis, which largely generates insertion and deletions (**Li et al., 2017**). In contrast, the EMS-induced mutations are mainly transitions, while transversions, InDels, and insertions are less frequent. In essence, the spectrum of mutations generated by EMS and genome editing are widely different. While genome-editing generates mainly null or amorphic mutants, the EMS-mutagenesis provides a broader range encompassing amorphic, hypomorphic, hypermorphic, and antimorphic mutants. The CRISPR/Cas9-based mutagenesis essentially extends the repertoire of mutations in crop species by providing additional variants not generated by EMS. Notwithstanding the above, our fully sequenced mutant collections allow gene functions investigation without the rigmarole of transformation and resources needed for genome editing.

## Conclusion

To sum up, our gene-indexed genome-wide mutant repertoire provides a resource to the scientific community to functionally characterize a gene or a set of the genes, including the unannotated genes. Our mutant resource can be used for improving a wide range of traits such as disease resistance, abiotic stress resistance and fruit-ripening, and basic studies in functional genomics. Our data is available through the tomato genome database-ITGV, where users can search for mutations in a desired gene or promoter and request the mutant seeds.

## Materials and methods

### Mutant population and DNA isolation

The doubly mutagenized (120 mM EMS) tomato (cultivar Arka Vikas) lines used in this study were the M_4_ progeny of the M_2_M_2_ population described earlier by **Gupta et al. (2017).** The M_2_M_2_ generation was carried forward to M_2_M_4_ plants. Leaf samples for genomic DNA were collected from 132 randomly selected M_2_M_4_ plants. Genomic DNA was isolated using DNeasy Plant Mini Kit (Qiagen) and in-house lab protocol (**Gupta et al., 2020b, Sreelakshmi et al., 2010**).

### DNA Sequencing, Read Mapping, and Variant Calling

Whole Genome Sequencing was performed on the HiSeqX sequencing system (Illumina) by GeneWiz Inc., NJ, following the manufacturer’s protocol. For each mutant line, a minimum of ∼200 million reads (30 Gb/sample) with a target of a minimum of 25 Gb > Q30 were generated. The raw reads were filtered using fastp software (v0.19.5) using parameters -M 30 -3 -5. The 2X 150 bp reads were mapped on *S. lycopersicum* cv. Heinz version SL3.0 using BWA-MEM (0.7.17) (**Li and Durbin, 2009**). The data analysis and variant calling was performed as described in **Gupta et al. (2020).**

Briefly, GATK (4.0.3.0) was used for generating BAM files, PCR duplicates removal, and variant calling. Variant filtration was carried out using the GATK VariantFiltration command. The parameters used for the filtering for SNPs were QualByDepth (QD <2), FisherStrand (FS>60), RMSMappingQuality (MQ<40), MQRankSum (−12.5), and ReadPosRankSum (−8.0); and for INDELs were QualByDepth (QD<2), FisherStrand (FS>200) and ReadPosRankSum (−20.0) (https://gatk.broadinstitute.org/hc/en-us/articles/360035531112?id=6925). The resulting vcf (variant calling format) files were annotated using the SIFT4G algorithm (Vaser et al., 2016). To validate the mutations identified, we selected 98 SNPs (71 heterozygous and 27 homozygous). These SNPs were subjected to Sanger sequencing, and all were confirmed positive (**Table S15**).

The effect of base substitutions on protein function was determined by SIFT4G (SIFT score ≤0.05 is considered deleterious) using the SIFT4G-ITAG3.2 genome reference database generated by Gupta et al. (2020). The deleteriously predicted genes by SIFT4G (≤0.05) were mapped on the tomato metabolic pathway (ITAG annotation 3.2; PlantCYC database 5.0.1) (Schläpfer et al., 2017). The genome-wide distribution of SNPs was generated using CircosVCF (Drori et al., 2017). Tomato microRNA loci coordinates were from the PmiREN database (https://www.pmiren.com/) (Guo et al., 2020). The amino acid circular plot was generated using the chorddiag package in R (https://github.com/mattflor/chorddiag).

### Carotenoid analysis and chloroplast movement

Carotenoid extraction and analysis was carried out as described by **Gupta et al. (2015)**. Chloroplast movement in tomato leaf was monitored by measuring the red light transmittance through leaf discs using a microplate reader (Biotek, Synergy HT) at 25°C as described by **Kilambi et al. (2021).**

### ITGV Database

The Induced Tomato Genomic Variant Database (ITGV) was made as described previously in **Gupta et al. (2020).** Briefly, the database runs on the XAMPP Apache server. The database was developed using MySQL/MariaDB relational database management system, HTML, CSS, PHP, and JavaScript libraries. Data search and submission queries were built using SQL. The genome browser ‘JBrowse’ was also integrated into the ITGV database to visualize the variants. The VCF files for all the accessions can be downloaded from the download data option in the database. The interested researchers can search ITGV online and identify mutations along with the SIFT scores in their target gene(s). The seeds of the mutant lines can be requested by using the online form provided on the ITGV website (http://psd.uohyd.ac.in/itgv/).

## Author Contributions

PG, YS, and RS designed this project and wrote the manuscript. PG performed most of the experiments. PSD made the database and wrote the code for it. KP and ASM characterized phototropin 1 mutants. All authors read and approved the manuscript.

## Acknowledgments

This work was supported by the Department of Biotechnology (DBT), India grants, BT/PR11671/PBD/16/828/2008, BT/PR/7002/PBD/16/1009/2012, and BT/COE/34/SP15209/2015 to RS and YS, and BT/PR6983/PBD/16/1007/2012,

BT/INF/22/SP44787/2021 to YS and RS. PSD acknowledge Bioinformatics Infrastructure Facility (BIF) at School of Life Sciences, University of Hyderabad.

## Supporting Information: A

**Figure S1.** EMS-induced nucleotide mispairing leading to GC>AT and AT>GC transitions.

**Figure S2.** Pie charts showing the frequency of transitions and transversions.

**Figure S3.** The total number of unique SNPs present in mutant lines in comparison to wild relatives and cultivars.

**Figure S4.** The ratio between the frequency of a mutated codon and its normal frequency in tomato.

**Figure S5.** Distribution of mutations in Cellular component and Molecular function GO categories.

**Figure S6.** Folate biosynthesis pathway marked with deleterious mutations present in the mutagenized population.

**Figure S7.** Light-signaling pathway marked with deleterious mutations present in the mutagenized population

**Figure S8.** The chloroplast relocation response in leaves of mutant line 142, and their backcrossed progeny.

**Figure S9.** The theoretically expected recovery of the recurrent parental line background on backcrossing.

**Table S1.** Summary of whole-genome sequencing of mutagenized lines.

**Table S2.** The percent nucleotide frequencies in +20 bp to -20 bp flanking the mutated base.

**Table S3.** Mutation density and distribution of SNPs in genic and intergenic regions of mutagenized lines.

**Table S4.** Chromosome-wise distribution of SNPs in mutagenized lines.

**Table S5.** Common and Unique SNPs present in mutagenized lines compared to tomato cultivars (**A**) and wild relatives (**B**).

**Table S6:** Distribution of different types of SNPs in the CDS of mutagenized lines.

**Table S7:** Distribution of INDELs in the CDS region of mutagenized lines.

**Table S8:** List of microRNAs regions bearing mutations in mutagenized lines.

**Table S9:** Amino acid change frequency in the mutagenized population.

**Table S10:** The frequency of individual mutated codons in the mutagenized population compared with codon usage frequency in tomato.

**Table S11:** The frequency of individual SNPs in natural accessions compared with codon usage frequency in tomato.

**Table S12:** The frequency of nonsynonymous mutations for individual codons compared with the theoretically predicted frequency.

**Table S13:** Classification of mutated genes in different gene ontology categories.

**Table S14:** Revalidation of mutations by Sanger sequencing.

**Table S15.** List of deleteriously predicted gene id’s in the whole mutant population

## References

1. Addo-Quaye C, Buescher E, Best N, Chaikam V, Baxter I, Dilkes BP (2017) Forward genetics by sequencing EMS variation-induced inbred lines. G3: Genes, Genomes, Genetics 7: 413–425

2. Aflitos S, Schijlen E, de Jong H, de Ridder D, Smit S, Finkers R, Wang J, Zhang G, Li N, Mao L et al. (2014) Exploring genetic variation in the tomato (Solanum section Lycopersicon) clade by wholeLJgenome sequencing. The Plant Journal 80: 136–148

3. Ashburner M, Golic KG, Hawley RS (2004) Drosophila: a laboratory handbook. Cold Spring Harbor Laboratory Press

4. Bai M, Yuan J, Kuang H, Gong P, Li S, Zhang Z, Liu B, Sun J, Yang M, Yang L et al. (2020) Generation of a multiplex mutagenesis population via pooled CRISPR-Cas9 in soybean. Plant Biotechnology Journal 18: 721–731

5. Baranczewski P, Nehls P, Rieger R, Pich U, Rajewsky MF, Schubert I (1997a) Formation and repair of O6-methylguanine in recombination hot spots of plant chromosomes. Environmental and Molecular Mutagenesis 29: 394–399

6. Baranczewski P, Nehls P, Rieger R, Rajewsky MF, Schubert I (1997b) Removal of O6-methylguanine from plant DNA in vivo is accelerated under conditions of clastogenic adaptation. Environmental and Molecular Mutagenesis 29: 400–405

7. Bauchet G, Causse M (2012) Genetic diversity in tomato (Solanum lycopersicum) and its wild relatives. Genetic Diversity in Plants 8: 134–162

8. Belkadi A, Bolze A, Itan Y, Cobat A, Vincent QB, Antipenko A, Shang L, Boisson B, Casanova J-L, Abel L (2015) Whole-genome sequencing is more powerful than whole-exome sequencing for detecting exome variants. Proceedings of the National Academy of Sciences 112: 5473–5478

9. Bulmer M (1991) The selection-mutation-drift theory of synonymous codon usage. Genetics 129: 897–907

10. Chen S, Zhou Y, Chen Y, Gu J (2018) fastp: an ultra-fast all-in-one FASTQ preprocessor. Bioinformatics 34: i884–i890

11. Comai L, Young K, Till BJ, Reynolds SH, Greene EA, Codomo CA, Enns LC, Johnson JE, Burtner C, Odden AR, et al. (2004) Efficient discovery of DNA polymorphisms in natural populations by Ecotilling. The Plant Journal 37: 778–786

12. Dickinson ME, Flenniken AM, Ji X, Teboul L, Wong MD, White JK, Meehan TF, Weninger WJ, Westerberg H, Adissu H, Baker CN et al. (2016) High-throughput discovery of novel developmental phenotypes. Nature 537: 508–514

13. Drake JW, Baltz RH (1976) The biochemistry of mutagenesis. Annual Review of Biochemistry 45: 11–37

14. Drori E, Levy D, Smirin-Yosef P, Rahimi O, Salmon-Divon M (2017) CircosVCF: circos visualization of whole-genome sequence variations stored in VCF files. Bioinformatics 33: 1392–1393

15. Fantini E, Falcone G, Frusciante S, Giliberto L, Giuliano G (2013) Dissection of tomato lycopene biosynthesis through virus-induced gene silencing. Plant Physiology 163: 986–998

16. Farrell A, Coleman BI, Benenati B, Brown KM, Blader IJ, Marth GT, Gubbels M-J (2014) Whole genome profiling of spontaneous and chemically induced mutations in Toxoplasma gondii. BMC Genomics 15: 1–15

17. Foolad MR (2007) Genome mapping and molecular breeding of tomato. International Journal of Plant Genomics. DOI: 10.1155/2007/64358

18. Gady AL, Hermans FW, Van de Wal MH, van Loo EN, Visser RG, Bachem CW (2009) Implementation of two high through-put techniques in a novel application: detecting point mutations in large EMS mutated plant populations. Plant Methods 5: 1–14

19. Garcia V, Bres C, Just D, Fernandez L, Tai FWJ, Mauxion J-P, Le Paslier M-C, Berard A, Brunel D, Aoki K. et al. (2016) Rapid identification of causal mutations in tomato EMS populations via mapping-by-sequencing. Nature Protocols 11: 2401–2418

20. Greene EA, Codomo CA, Taylor NE, Henikoff JG, Till BJ, Reynolds SH, Enns LC, Burtner C, Johnson JE, Odden AR et al. (2003) Spectrum of chemically induced mutations from a large-scale reverse-genetic screen in Arabidopsis. Genetics 164: 731–740

21. Guo Y, Abernathy B, Zeng Y, Ozias-Akins P (2015) TILLING by sequencing to identify induced mutations in stress resistance genes of peanut (Arachis hypogaea). BMC Genomics 16: 1–13

22. Guo Z, Kuang Z, Wang Y, Zhao Y, Tao Y, Cheng C, Yang J, Lu X, Hao C, Wang T et al. (2020) PmiREN: a comprehensive encyclopedia of plant miRNAs. Nucleic acids research 48: D1114–D1121

23. Gupta P, Dholaniya PS, Devulapalli S, Tawari NR, Sreelakshmi Y, Sharma R (2020a) Reanalysis of genome sequences of tomato accessions and its wild relatives: development of Tomato Genomic Variation (TGV) database integrating SNPs and INDELs polymorphisms. Bioinformatics 36: 4984–4990

24. Gupta P, Reddaiah B, Salava H, Upadhyaya P, Tyagi K, Sarma S, Datta S, Malhotra B, Thomas S, Sunkum A et al. (2017) Next-generation sequencing (NGS)-based identification of induced mutations in a doubly mutagenized tomato (Solanum lycopersicum) population. The Plant Journal 92: 495–508

25. Gupta P, Salava H, Sreelakshmi Y, Sharma R (2020b) A low-cost high-throughput method for plant genomic DNA isolation. In Cereal Genomics. Springer, pp 1–7

26. Gupta P, Sreelakshmi Y, Sharma R (2015) A rapid and sensitive method for determination of carotenoids in plant tissues by high performance liquid chromatography. Plant Methods 11: 1–12

27. Henry IM, Nagalakshmi U, Lieberman MC, Ngo KJ, Krasileva KV, Vasquez-Gross H, Akhunova A, Akhunov E, Dubcovsky J, Tai TH et al. (2014) Efficient genome-wide detection and cataloging of EMS-induced mutations using exome capture and next-generation sequencing. The Plant Cell 26: 1382–1397

28. Hershberg R, Petrov DA (2008) Selection on codon bias. Annual Review of Genetics 42: 287–299

29. Hershberg R, Petrov DA (2009) General rules for optimal codon choice. PLoS Genetics 5: e1000556

30. Hospital F (2003) Marker-assisted breeding. In: H.J. Newbury, editor. Plant molecular breeding. Oxford and Boca Raton: Blackwell Publishing and CRC Press, 30–59

31. Ikemura T (1985) Codon usage and tRNA content in unicellular and multicellular organisms. Molecular Biology and Evolution 2: 13–34

32. Jacob P, Avni A, Bendahmane A (2018) Translational research: exploring and creating genetic diversity. Trends in Plant Science 23: 42–52

33. Kilambi HV, Dindu A, Sharma K, Nizampatnam NR, Gupta N, Thazath NP, Dhanya AJ, Tyagi K, Sharma S, Sharma R et al. (2021) The new kid on the block: A dominant-negative mutation of phototropin1 enhances carotenoid content in tomato fruits. The Plant Journal 106: 844–861

34. Krasileva KV, Vasquez-Gross HA, Howell T, Bailey P, Paraiso F, Clissold L, Simmonds J, Ramirez-Gonzalez RH, Wang X, Borrill P. et al. (2017) Uncovering hidden variation in polyploid wheat. Proceedings of the National Academy of Sciences 114: E913–E921

35. Kulus D (2018) Genetic resources and selected conservation methods of tomato. Journal of Applied Botany and Food Quality 91, 135–144

36. Kumar P, Henikoff S, Ng PC (2009) Predicting the effects of coding non-synonymous variants on protein function using the SIFT algorithm. Nature Protocols 4: 1073–1081

37. Leitao J, Shu Q, Forster B, Nakagawa H (2011) Chemical mutagenesis,” in Plant Mutation Breeding and Biotechnology Joint FAO/IAEA Programme, Eds B. P. Forster, Q. Y. Shu, and H. Nakagawa (Wallingford: CABI), 135–158.

38. Li G, Jain R, Chern M, Pham NT, Martin JA, Wei T, Schackwitz WS, Lipzen AM, Duong PQ, Jones KC et al. (2017) The sequences of 1504 mutants in the model rice variety Kitaake facilitate rapid functional genomic studies. The Plant Cell 29: 1218–1231

39. Li H, Durbin R (2009) Fast and accurate short read alignment with Burrows–Wheeler transform. Bioinformatics 25: 1754–1760

40. Lin T, Zhu G, Zhang J, Xu X, Yu Q, Zheng Z, Zhang Z, Lun Y, Li S, Wang X et al. (2014) Genomic analyses provide insights into the history of tomato breeding. Nature Genetics 46: 1220–1226

41. Liu H-J, Jian L, Xu J, Zhang Q, Zhang M, Jin M, Peng Y, Yan J, Han B, Liu J,., et al. (2020) High-throughput CRISPR/Cas9 mutagenesis streamlines trait gene identification in maize. The Plant Cell 32: 1397–1413

42. Lu X, Liu J, Ren W, Yang Q, Chai Z, Chen R, Wang L, Zhao J, Lang Z, Wang H. et al., (2018) Gene-indexed mutations in maize. Molecular Plant 11: 496–504

43. Lu Y, Ye X, Guo R, Huang J, Wang W, Tang J, Tan L, Zhu J-K, Chu C, Qian Y. (2017) Genome-wide targeted mutagenesis in rice using the CRISPR/Cas9 system. Molecular Plant 10: 1242–1245

44. Lyons DM, Lauring AS (2017) Evidence for the selective basis of transition-to-transversion substitution bias in two RNA viruses. Molecular Biology and Evolution 34: 3205–3215

45. Ma L, Mu J, Grierson D, Wang Y, Gao L, Zhao X, Zhu B, Luo Y, Shi K, Wang Q. et al. (2020) Noncoding RNAs: functional regulatory factors in tomato fruit ripening. Theoretical and Applied Genetics 133: 1753–1762

46. Manova V, Gruszka D (2015) DNA damage and repair in plants–from models to crops. Frontiers in Plant Science 6: 885

47. Marroni F, Pinosio S, Di Centa E, Jurman I, Boerjan W, Felice N, Cattonaro F, Morgante M (2011) Large-scale detection of rare variants via pooled multiplexed next-generation sequencing: towards next-generation Ecotilling. The Plant Journal 67: 736–745

48. McCallum CM, Comai L, Greene EA, Henikoff S (2000) Targeting Induced Local Lesions IN Genomes (TILLING) for plant functional genomics. Plant Physiology 123: 439–442

49. Menda N, Semel Y, Peled D, Eshed Y, Zamir D (2004) In silico screening of a saturated mutation library of tomato. The Plant Journal 38: 861–872

50. Meng X, Yu H, Zhang Y, Zhuang F, Song X, Gao S, Gao C, Li J (2017) Construction of a genome-wide mutant library in rice using CRISPR/Cas9. Molecular Plant 10: 1238–1241

51. Minoia S, Petrozza A, D’Onofrio O, Piron F, Mosca G, Sozio G, Cellini F, Bendahmane A, Carriero F (2010) A new mutant genetic resource for tomato crop improvement by TILLING technology. BMC Research Notes 3: 1–8

52. Monroe J, Srikant T, Carbonell-Bejerano P, Becker C, Lensink M, Exposito-Alonso M, Klein M, Hildebrandt J, Neumann M, Kliebenstein D. et al. (2022) Mutation bias reflects natural selection in Arabidopsis thaliana. Nature 602: 101–105.

53. Ng PC, Henikoff S (2003) SIFT: Predicting amino acid changes that affect protein function. Nucleic Acids Research 31: 3812–3814

54. Okabe Y, Ariizumi T (2016) Mutant resources and TILLING platforms in tomato research. *In* Functional Genomics and Biotechnology in Solanaceae and Cucurbitaceae crops. Eds. i Ezura E, Ariizumi T, Garcia-Mas J, Rose J, Editors Springer, pp 75–91

55. Okabe Y, Ariizumi T, Ezura H (2013) Updating the micro-tom TILLING platform. Breeding Science 63: 42–48

56. Okabe Y, Asamizu E, Saito T, Matsukura C, Ariizumi T, Bres C, Rothan C, Mizoguchi T, Ezura H (2011) Tomato TILLING technology: development of a reverse genetics tool for the efficient isolation of mutants from Micro-Tom mutant libraries. Plant and Cell Physiology 52: 1994–2005

57. Oleykowski CA, Bronson Mullins CR, Godwin AK, Yeung AT (1998) Mutation detection using a novel plant endonuclease. Nucleic Acids Research 26: 4597–4602

58. Pegg AE (2011) Multifaceted roles of alkyltransferase and related proteins in DNA repair, DNA damage, resistance to chemotherapy, and research tools. Chemical Research in Toxicology 24: 618–639

59. Piron F, Nicolaï M, Minoïa S, Piednoir E, Moretti A, Salgues A, Zamir D, Caranta C, Bendahmane A (2010) An induced mutation in tomato eIF4E leads to immunity to two potyviruses. PloS one 5: e11313

60. Provart NJ, Alonso J, Assmann SM, Bergmann D, Brady SM, Brkljacic J, Browse J, Chapple C, Colot V, Cutler S,., et al., (2016) 50 years of Arabidopsis research: highlights and future directions. New Phytologist 209: 921–944

61. Ram H, Soni P, Salvi P, Gandass N, Sharma A, Kaur A, Sharma TR (2019) Insertional mutagenesis approaches and their use in rice for functional genomics. Plants 8: 310

62. Rigola D, van Oeveren J, Janssen A, Bonné A, Schneiders H, van der Poel HJ, van Orsouw NJ, Hogers RC, de Both MT, van Eijk MJ (2009) High-throughput detection of induced mutations and natural variation using KeyPoint™ technology. PloS one 4: e4761

63. Schläpfer P, Zhang P, Wang C, Kim T, Banf M, Chae L, Dreher K, Chavali AK, Nilo-Poyanco R, Bernard T. et al. (2017) Genome-wide prediction of metabolic enzymes, pathways, and gene clusters in plants. Plant Physiology 173: 2041–2059

64. Sharma K, Gupta S, Sarma S, Rai M, Sreelakshmi Y, Sharma R (2021) Mutations in tomato 1-aminocyclopropane carboxylic acid synthase2 uncover its role in development beside fruit ripening. The Plant Journal 106: 95–112

65. Sharma S, Kharshiing E, Srinivas A, Zikihara K, Tokutomi S, Nagatani A, Fukayama H, Bodanapu R, Behera RK, Sreelakshmi Y. et al. (2014) A dominant mutation in the light-oxygen and voltage2 domain vicinity impairs phototropin1 signaling in tomato. Plant Physiology 164: 2030–2044

66. Shirasawa K, Hirakawa H, Nunome T, Tabata S, Isobe S (2016) Genome-wide survey of artificial mutations induced by ethyl methanesulfonate and gamma rays in tomato. Plant Biotechnology Journal 14: 51–60

67. Simko I, Jia M, Venkatesh J, Kang B-C, Weng Y, Barcaccia G, Lanteri S, Bhattarai G, Foolad MR (2021) Genomics and Marker-Assisted Improvement of Vegetable Crops. Critical Reviews in Plant Sciences 40: 303–365

68. Sreelakshmi Y, Gupta S, Bodanapu R, Chauhan VS, Hanjabam M, Thomas S, Mohan V, Sharma S, Srinivasan R, Sharma R (2010) NEATTILL: A simplified procedure for nucleic acid extraction from arrayed tissue for TILLING and other high-throughput reverse genetic applications. Plant Methods 6: 1–11

69. Stenøien HK (2005) Adaptive basis of codon usage in the haploid moss Physcomitrella patens. Heredity 94: 87–93

70. Tanksley SD, McCouch SR (1997) Seed banks and molecular maps: unlocking genetic potential from the wild. Science 277: 1063–1066

71. Thompson O, Edgley M, Strasbourger P, Flibotte S, Ewing B, Adair R, Au V, Chaudhry I, Fernando L, Hutter H. et al. (2013) The million mutation project: a new approach to genetics in Caenorhabditis elegans. Genome research 23: 1749–1762

72. Tieman D, Zhu G, Resende Jr MF, Lin T, Nguyen C, Bies D, Rambla JL, Beltran KSO, Taylor M, Zhang B (2017) A chemical genetic roadmap to improved tomato flavor. Science 355: 391–394

73. Till BJ, Cooper J, Tai TH, Colowit P, Greene EA, Henikoff S, Comai L (2007) Discovery of chemically induced mutations in rice by TILLING. BMC Plant Biology 7: 1–12

74. Tsai H, Howell T, Nitcher R, Missirian V, Watson B, Ngo KJ, Lieberman M, Fass J, Uauy C, Tran RK et al., (2011) Discovery of rare mutations in populations: TILLING by sequencing. Plant Physiology 156: 1257–1268

75. Tsai H, Missirian V, Ngo KJ, Tran RK, Chan SR, Sundaresan V, Comai L (2013) Production of a high-efficiency TILLING population through polyploidization. Plant Physiology 161: 1604–1614

76. Tsuda M, Kaga A, Anai T, Shimizu T, Sayama T, Takagi K, Machita K, Watanabe S, Nishimura M, Yamada N et al., (2015) Construction of a high-density mutant library in soybean and development of a mutant retrieval method using amplicon sequencing. BMC Genomics 16: 1–18

77. Uauy C, Paraiso F, Colasuonno P, Tran RK, Tsai H, Berardi S, Comai L, Dubcovsky J (2009) A modified TILLING approach to detect induced mutations in tetraploid and hexaploid wheat. BMC Plant Biology 9: 1–14

78. Vaser R, Adusumalli S, Leng SN, Sikic M, Ng PC (2016) SIFT missense predictions for genomes. Nature Protocols 11: 1–9

79. Wang P, Amato NJ, Zhai Q, Wang Y (2015) Cytotoxic and mutagenic properties of O^4^-alkylthymidine lesions in Escherichia coli cells. Nucleic Acids Research 43: 10795–10803

80. Wu J, Li L, Wang P, You C, Williams NL, Wang Y (2016) Translesion synthesis of O^4^-alkylthymidine lesions in human cells. Nucleic Acids Research 44: 9256–9265

81. Yan W, Deng XW, Yang C, Tang X (2021) The Genome-Wide EMS Mutagenesis Bias Correlates With Sequence Context and Chromatin Structure in Rice. Frontiers in Plant Science 12: 370

82. Yano R, Hoshikawa K, Okabe Y, Wang N, Dung PT, Imriani PS, Shiba H, Ariizumi T, Ezura H (2019) Multiplex exome sequencing reveals genome-wide frequency and distribution of mutations in the ‘Micro-Tom’ Targeting Induced Local Lesions In Genomes (TILLING) mutant library. Plant Biotechnology 36: 223–231

83. Zhang J (2000) Rates of conservative and radical nonsynonymous nucleotide substitutions in mammalian nuclear genes. Journal of Molecular Evolution 50: 56–68

84. Zhang M, Wang C, Otto TD, Oberstaller J, Liao X, Adapa SR, Udenze K, Bronner IF, Casandra D, Mayho M et al. (2018) Uncovering the essential genes of the human malaria parasite Plasmodium falciparum by saturation mutagenesis. Science 360: eaap7847

85. Zuo J, Grierson D, Courtney LT, Wang Y, Gao L, Zhao X, Zhu B, Luo Y, Wang Q, Giovannoni JJ (2020) Relationships between genome methylation, levels of non-coding RNAs, mRNAs and metabolites in ripening tomato fruit. The Plant Journal 103: 980–994

