## Supplementary material for "Whole-Genome Profiling of Ethyl Methanesulfonate Mutagenesis in Tomato": Figure S

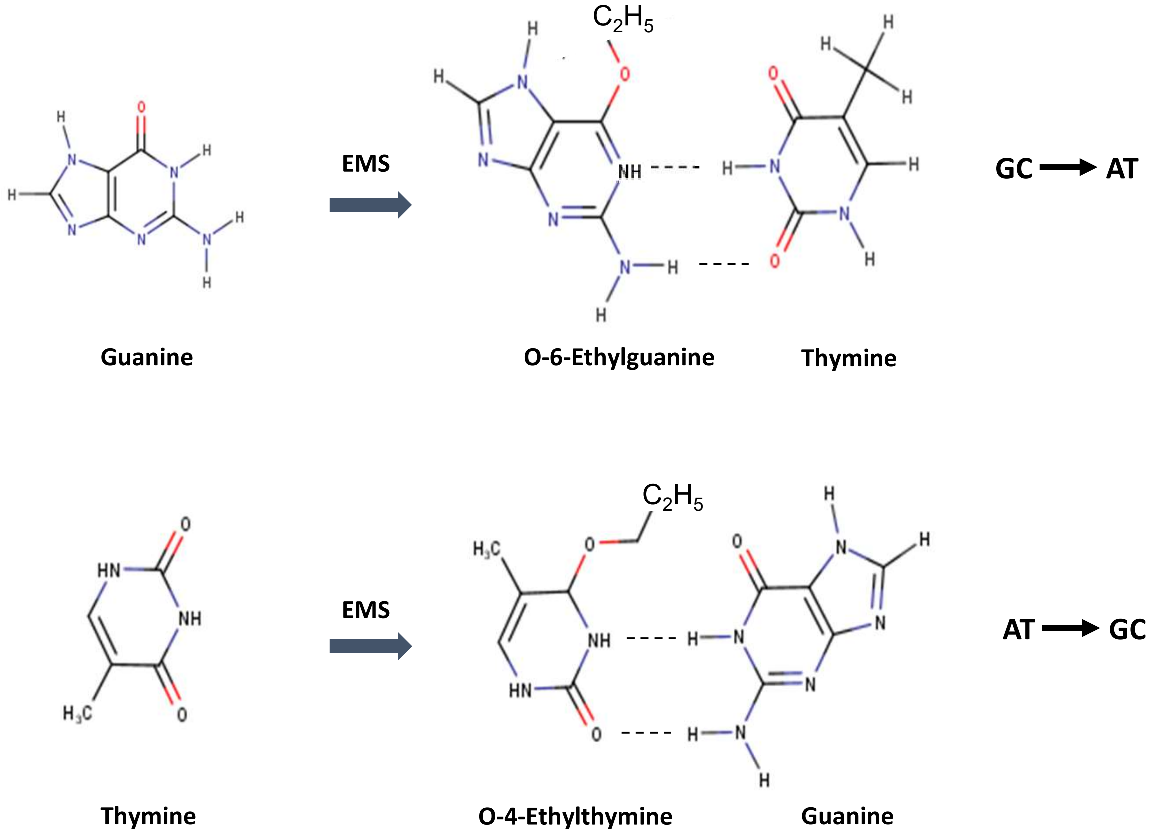


**Figure S1.** EMS-induced nucleotide mispairing leading to GC>AT and AT>GC transitions. EMS alkylates the $O^6$ position of guanine and $O^4$ position of thymine, leading to mispairing with thymine and guanine, respectively.

.

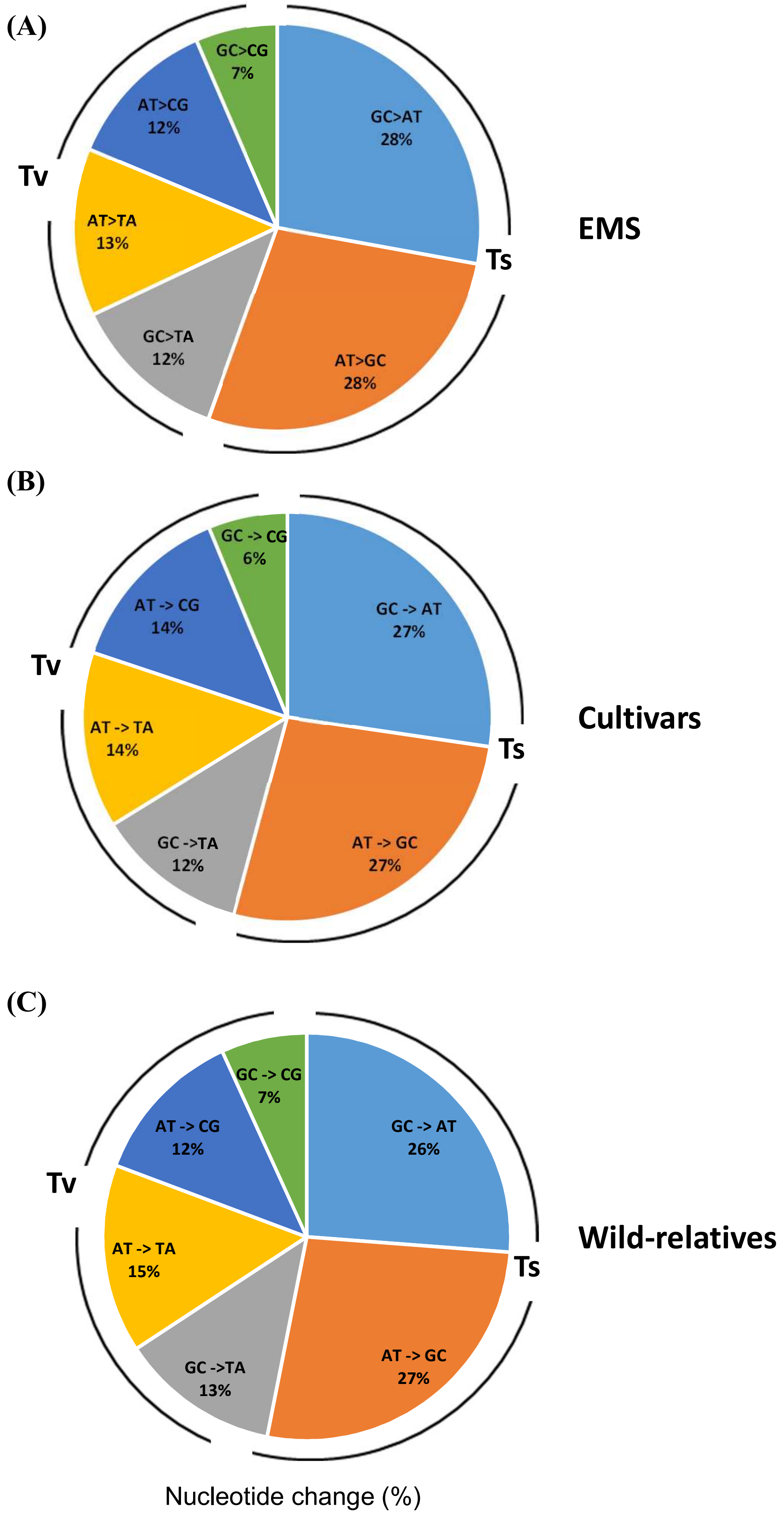


**Figure S2**. Pie charts showing the frequency of transitions and transversions. (**A**) EMS (Arka Vikas), (**B**) tomato cultivars, and (**C**) wild-relatives. **Ts**- transitions, **Tv**- transversions.

(A)

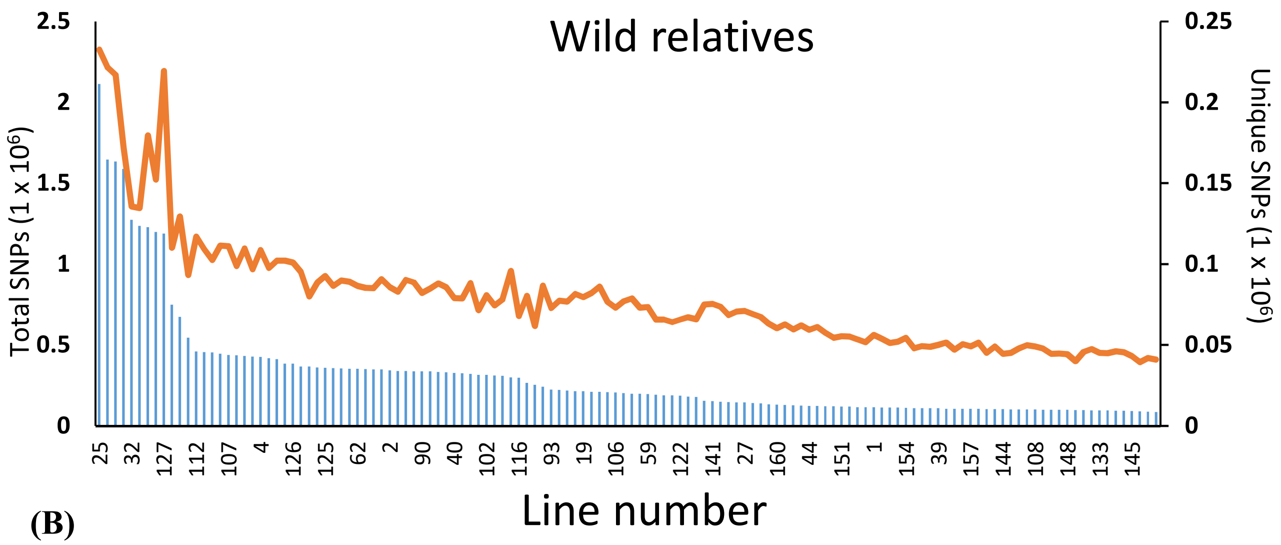


(B)

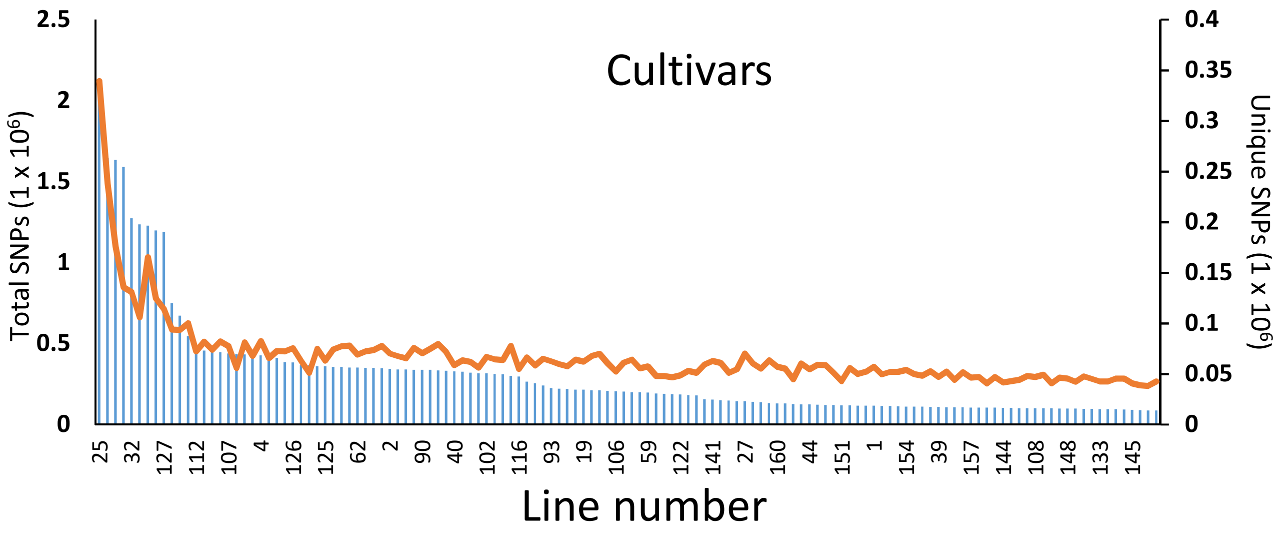


**Figure S3.** The total number of unique SNPs present in 132 mutant lines compared to 31 wild relatives of tomato (**A**) and 53 tomato cultivars (**B**). For details, see Table S5.

**(A)**

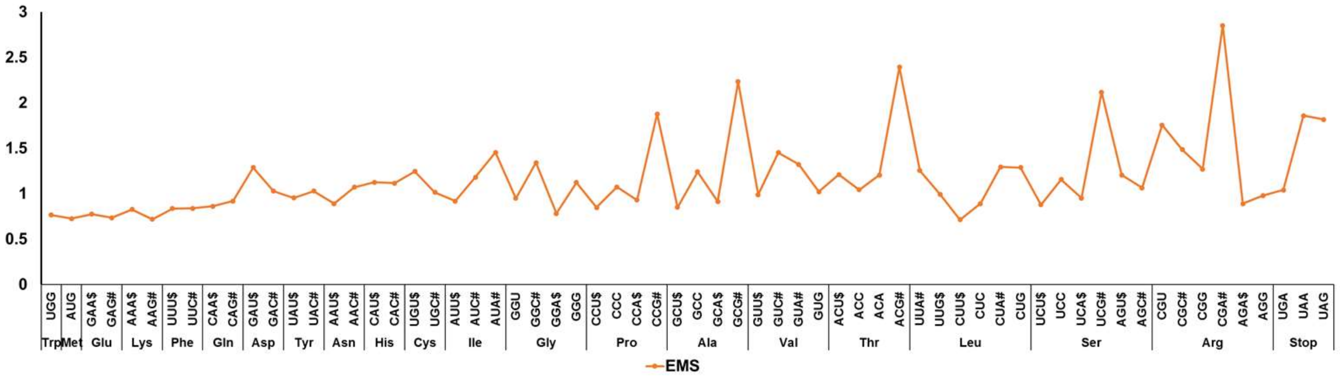


**(B)**

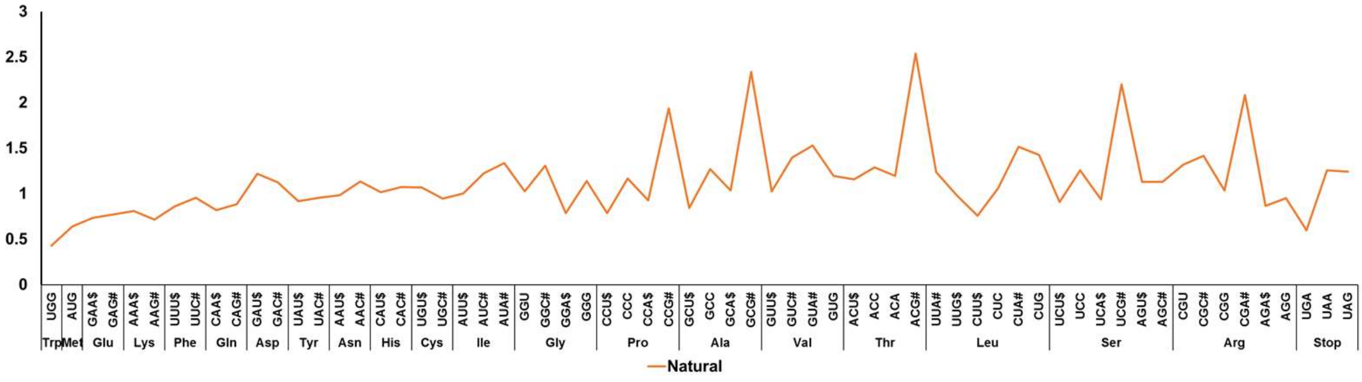


**(C)**

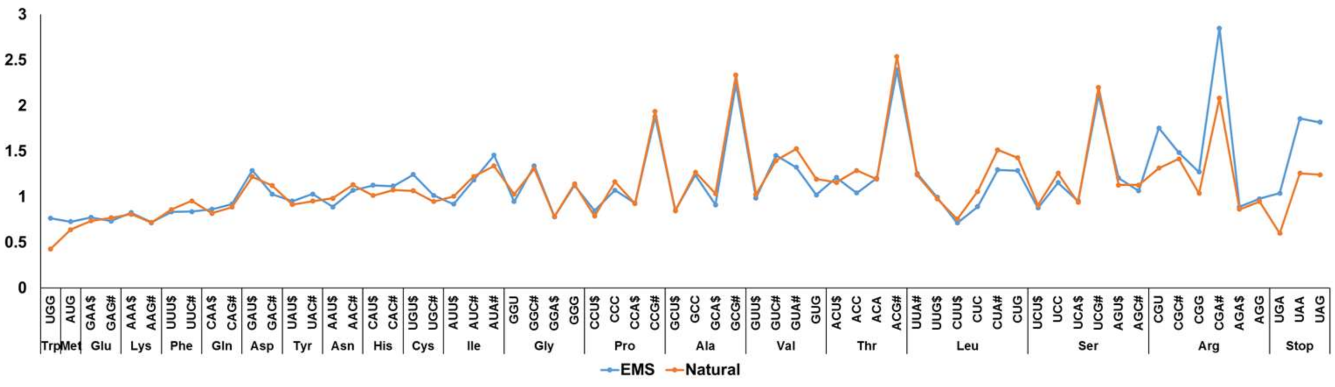


**Figure S4**. The ratio between the frequency of a mutated codon and its normal frequency in tomato. (**A**) EMS , (**B**) Natural and (**C**) EMS v/s Natural . The most preferred codon as per the tomato codon usage table (https://solgenomics.net/documents/misc/codon_usage/codon_usage_data/l_esculentum_codon_usage_table.txt) is marked with the **$** dollar sign, and least preferred codon is marked as # hashtag after the codon letter in the graph. The individual ratios are given in Table S11.

Cellular component

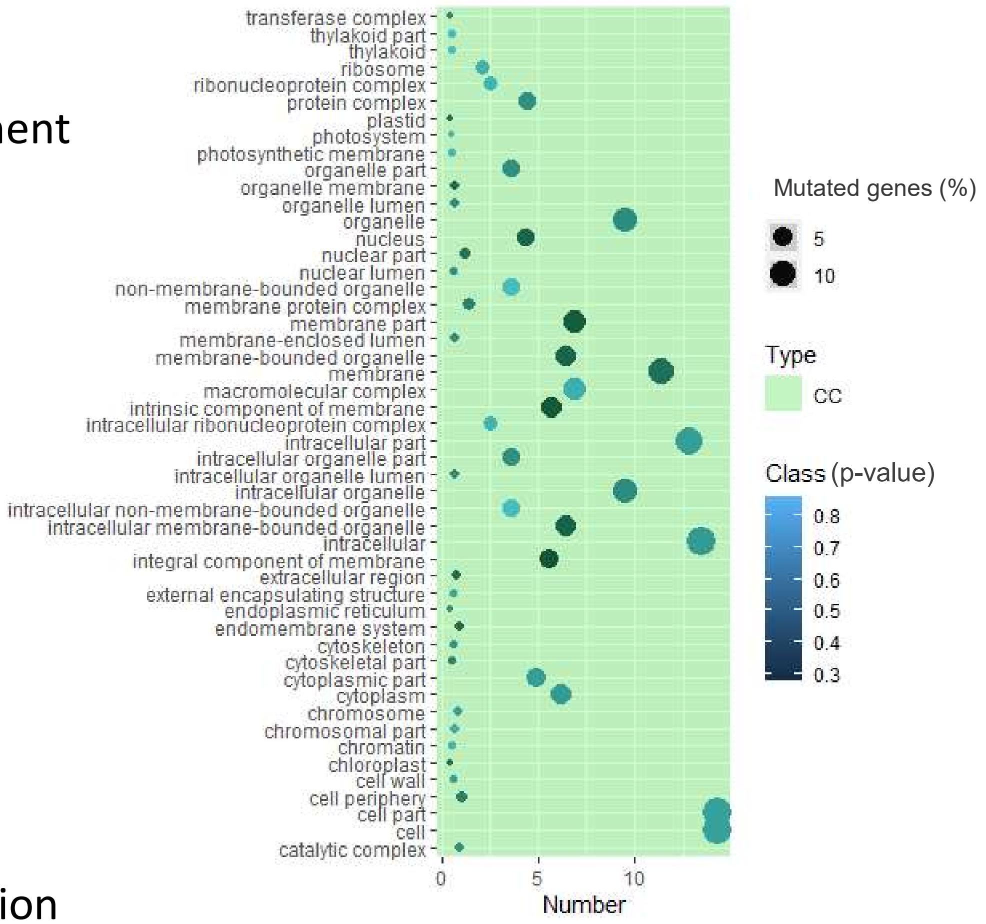


Molecular function

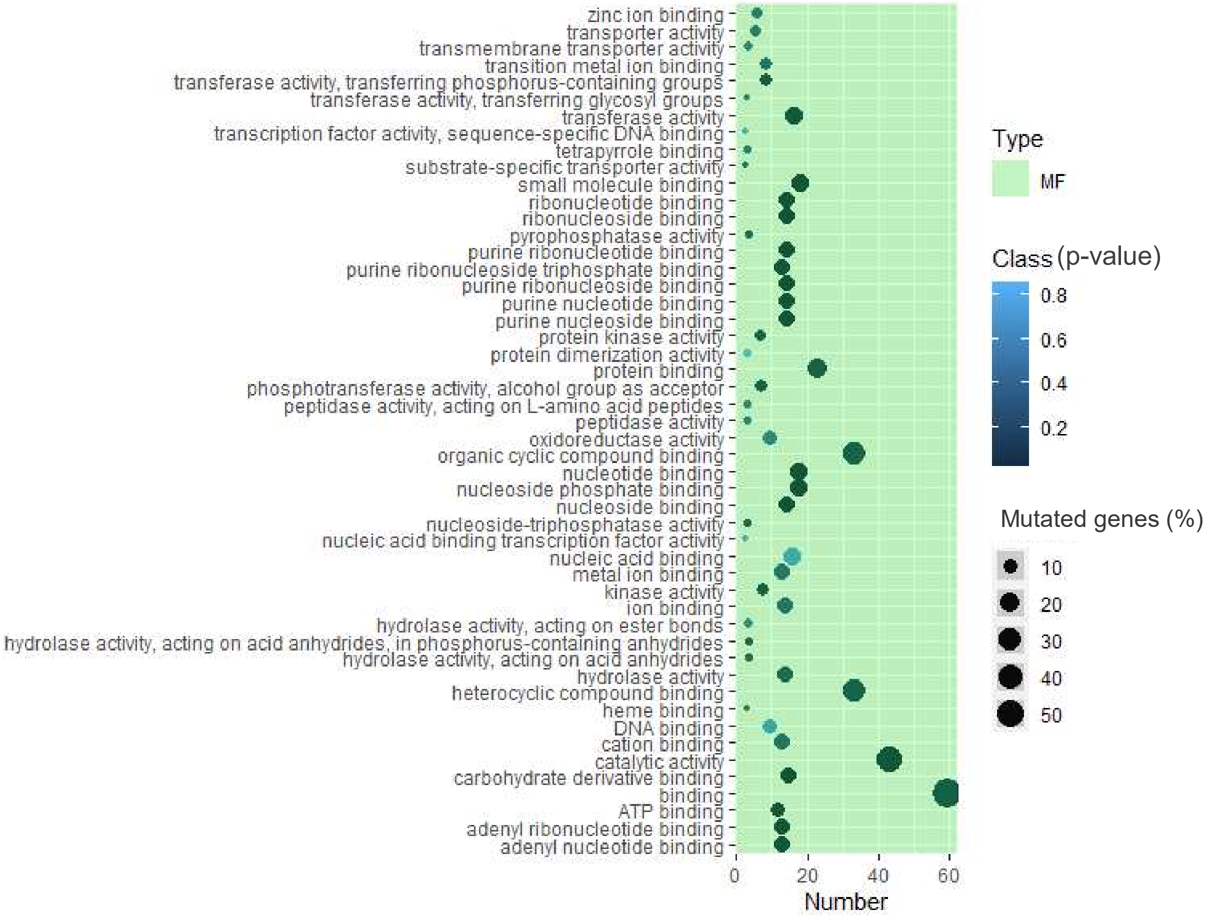


**Figure S5.** Distribution of mutations in Cellular component and Molecular function GO categories. Top 50 GO categories with the high frequency of mutations in Cellular component and Molecular function. For details, see Table S13.

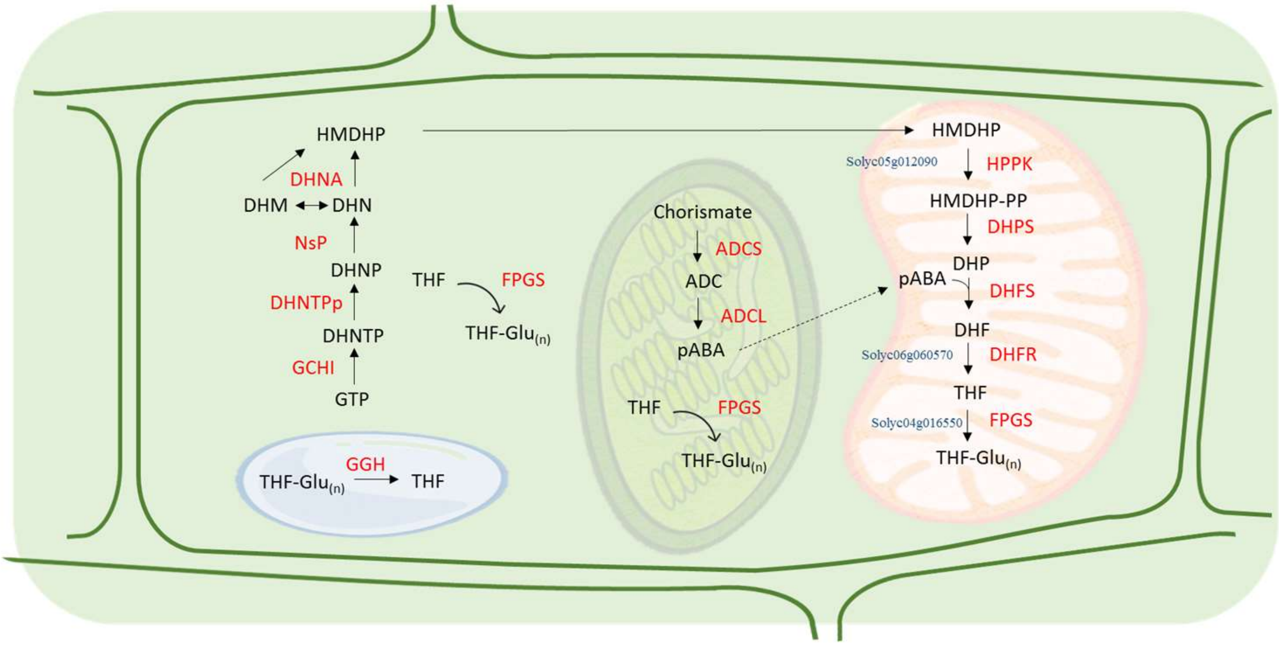


**Figure S6.** Folate biosynthesis pathway marked with deleterious mutations present in the mutagenized population. The gene id's are marked in blue on different steps of the pathway. Note that the folate biosynthetic pathway is recalcitrant to mutagenesis, as folate is essential for nucleotide synthesis and C1 metabolism. For details, see Table S15.

**Abbreviations:** *Precursors:* GTP, guanosine triphosphate; DHNTP, dihydroneopterin triphosphate; DHNP, dihydroneopterin monophosphate; DHN, dihydroneopterin; HMDHP, 6-hydroxymethyldihydropterin; HMDHP-PP, 6-hydroxymethyldihydropterin pyrophosphate; DHP, dihydropteroate; DHF, dihydrofolate; THF, tetrahydrofolate; THF-$Glu_{(n)}$, tetrahydrofolate polyglutamate; ADC, aminodeoxychorismate; pABA, para-aminobenzoic acid. *Enzymes:* GCHI, GTP cyclohydrolase I; $DHNTP_P$-diphosphatase, dihydroneopterin triphosphate pyrophosphatase; DHNA, dihydroneopterine aldolase; HPPK, HMDHP pyrophosphokinase; DHPS, dihydropteroate synthase; DHFR, dihydrofolate reductase; FPGS, folylpolyglutamate synthetase; ADCS, aminodeoxychorismate synthase; ADCL, aminodeoxychorismate lyase; GGH, gamma-glutamyl hydrolase.

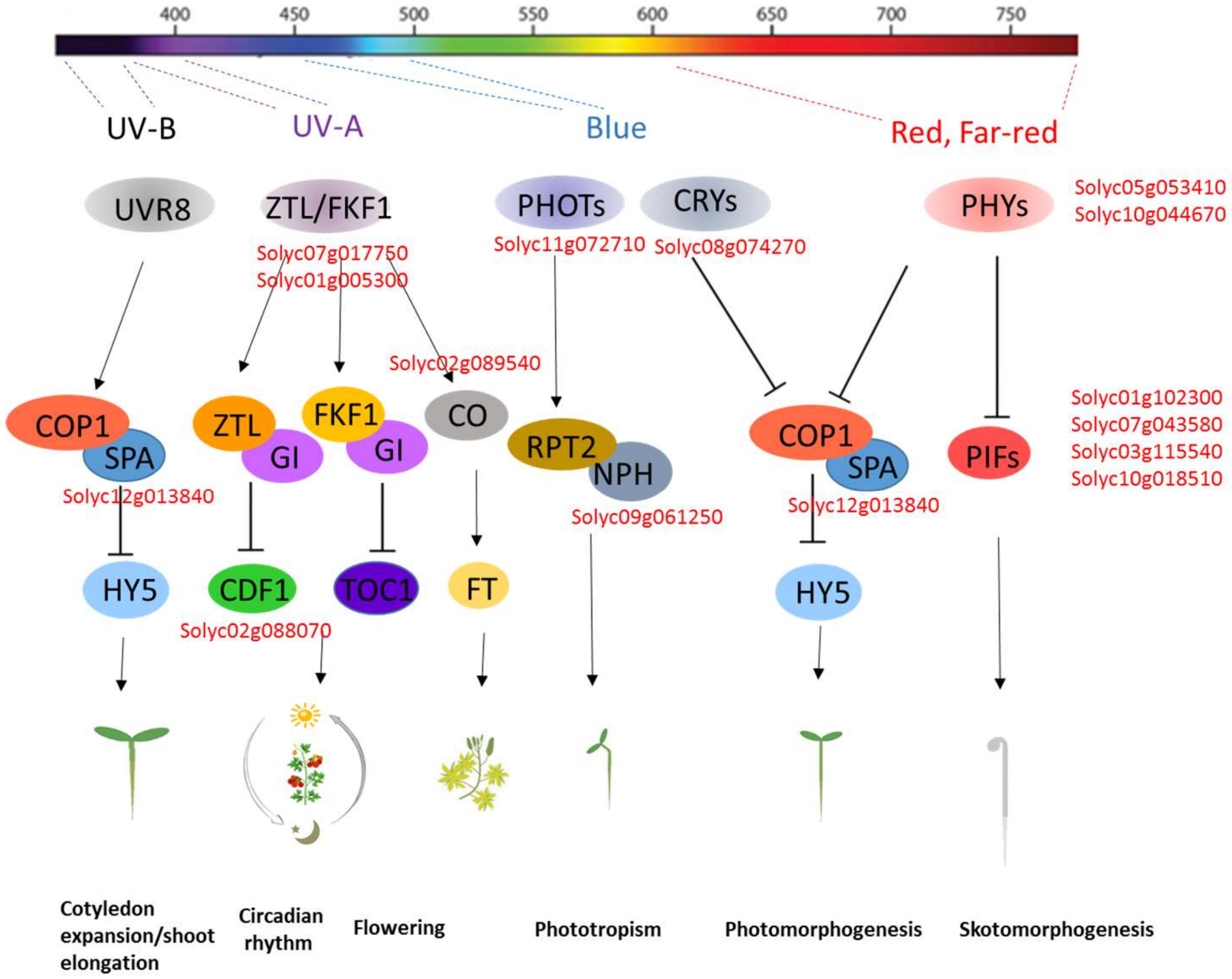


**Figure S7**. Light-signaling pathway marked with deleterious mutations present in the mutagenized population. The gene id's are marked in red on different steps of the pathway.

**Abbreviations:** UVR8, UV-B resistance 8; COP1, Constitutive Photomorphogenic 1; SPA, Sugar partitioning Affecting protein; HY5, Elongated Hypocotyl5; ZTL, ZEITLUPE; FKF1, Flavin-binding, Kelch Repeat, F-BOX 1; GI, GIGANTEA; CDF1, Cycling Dof Factor 1; TOC1, Timing Of Cab Expression 1; CO, CONSTANS; FT, Flowering Locus T; PHOTs, Phototropins; RPT2, Root Phototropism protein 2; NPH, Non-Phototropic Hypocotyl; CRYs, Cryptochromes; PHYs, Phytochromes; PIFs, Phytochrome interacting factors.

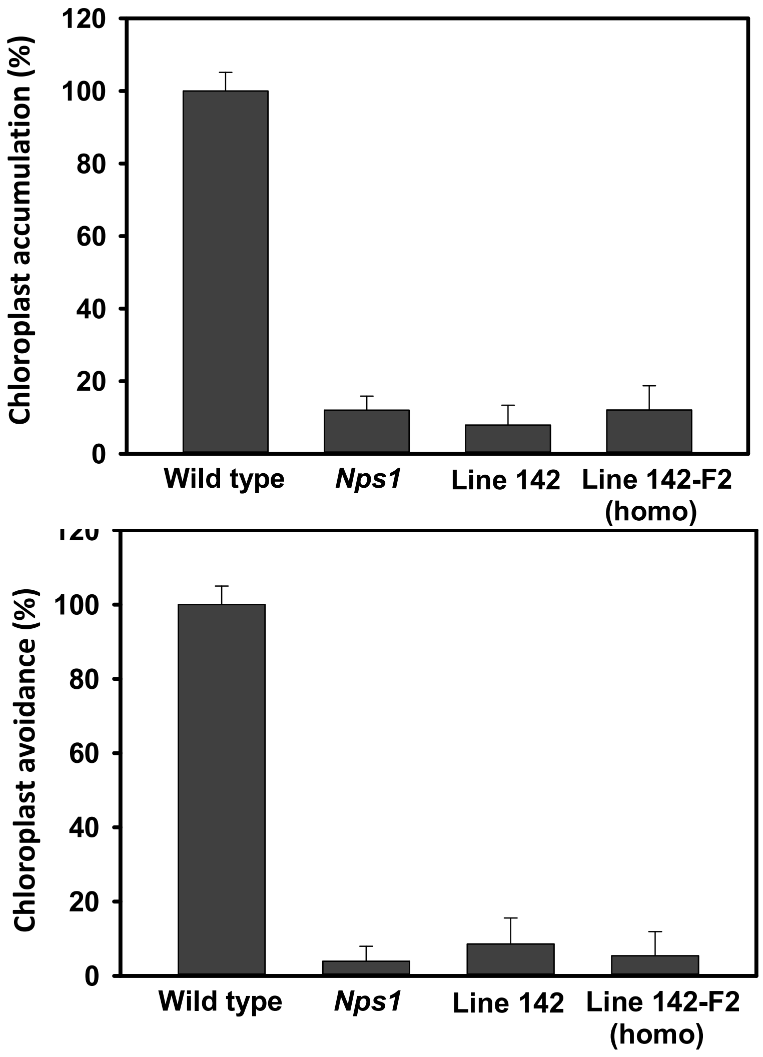


**Figure S8**. The chloroplast relocation response in leaves of mutant line 142, and their backcrossed progeny. Similar to the *Nps1* mutant, both lines show near-total loss of chloroplast relocation response in the parental line as well as in the $F_2$ generation.

.

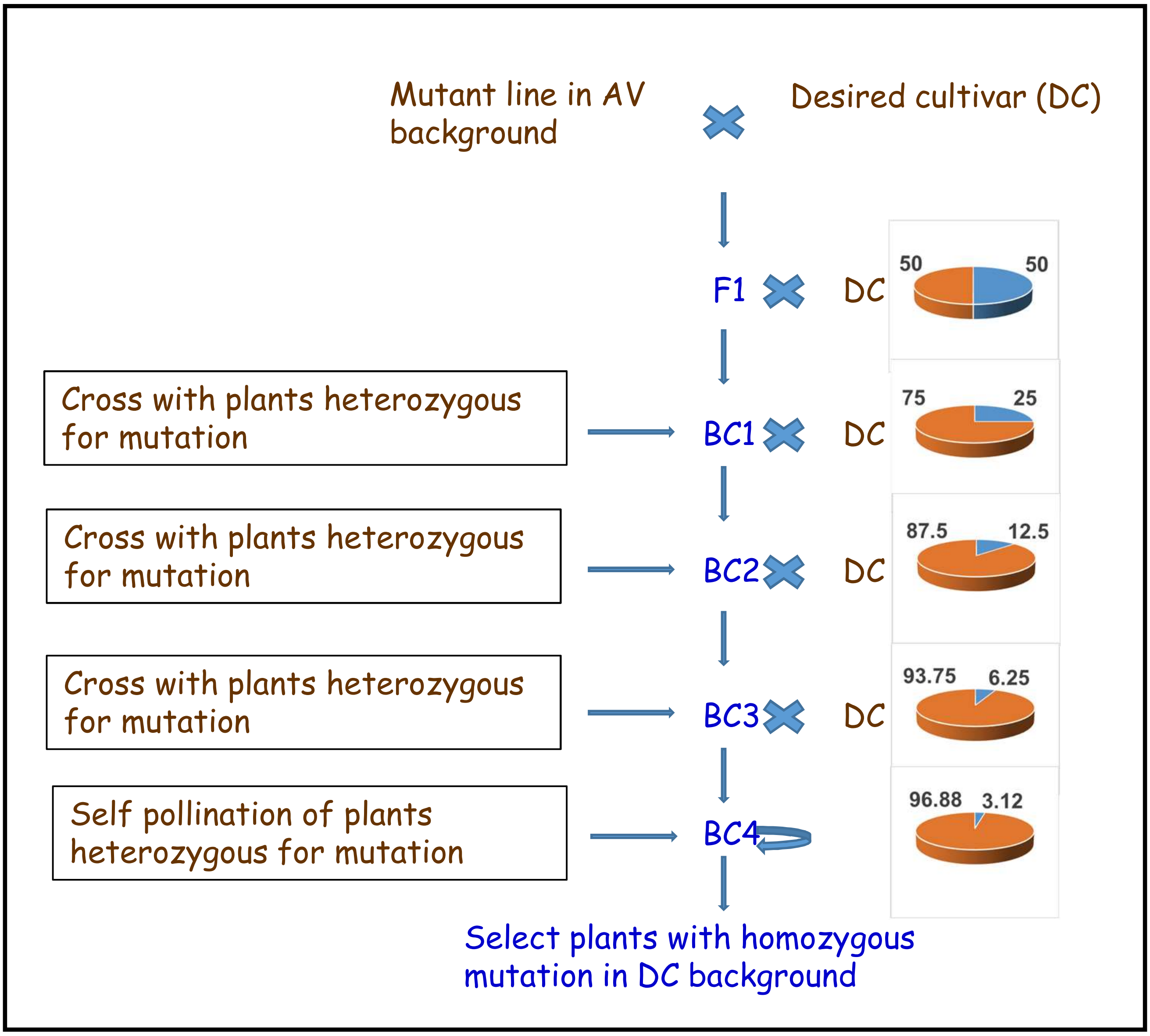


**Figure S9:** The theoretically expected recovery of the recurrent parental line background on backcrossing (red pie). Since the mutated gene itself acts as a marker, the expected recovery by $BC_4$ is ≥ 99% of the recurrent parental genome (**Hospital F**. 2003 Marker-assisted breeding**. In:** H.J. Newbury, editor. *Plant molecular breeding*. Oxford and Boca Raton: Blackwell Publishing and CRC Press; p. 30-59). By doing WGS of the $BC_4F_2$ lines homozygous for mutations, the near-isogenic lines to the recurrent parent can be recovered.
